# Mitochondria Controlled mTORC1 Activation Compartmentalizes Translation Initiation Factor eIF4E to Augment Intracellular Trafficking and Extracellular Export of miRNA in Mammalian Cells

**DOI:** 10.1101/2020.05.20.105601

**Authors:** Susanta Chatterjee, Yogaditya Chakrabarty, Saikat Banerjee, Souvik Ghosh, Suvendra N. Bhattacharyya

## Abstract

Defective intracellular trafficking and export of miRNAs has been observed in senescent mammalian cells having impaired mitochondrial potential. Similar to what happens in senescent cells, Uncoupling Protein 2 mediated depolarization of mitochondrial membrane potential results in progressive sequestration of miRNAs with polysomes and lowered release of miRNAs through extracellular vesicles. Supporting importance of mitochondrial membrane potential on miRNAs’ fate determination, impaired miRNA-trafficking process in growth retarded human cells has been found to be reversed in presence of Genipin an inhibitor of Uncoupling Protein 2. Mitochondrial detethering of endoplasmic reticulum in mitochondria depolarized cells, found to be responsible for defective compartmentalization of translation initiation factor eIF4E to ER attached polysomes. It causes retarded translation process of target mRNAs with rER attached polysomes to ensure reduced intracellular trafficking and extracellular export of miRNAs. We have identified a reduced activity of mTORC1 complex in mitochondria defective cells to cause reduced phosphorylation of eIF4E-BP1 to cause retarded eIF-4E targeting to ER attached polysome. Cumulatively, these data suggest intricate involvement of mitochondrial membrane potential and dynamics to determine stability of miRNAs in mammalian cells by affecting sub-cellular locations and export of miRNPs by affecting mTORC1 complex, the regulator of the protein translational machinery.

**Significance statement:** How the reduced mitochondrial activity in growth retarded cells causes defective miRNA export is an open question. Mitochondrial defects induces a retarded subcellular miRNP trafficking in human cells to cause an upregulation in cellular miRNA content by reducing extracellular vesicle-mediated export of miRNA. We have identified a defective compartmentalization of translation initiation factor eIF4E in mitochondria-ER detethered mammalian cells to cause the retarded intracellular miRNA movement and export Activity of mTORC1 complex, a key regulator of protein translation in mammalian cells, is found to be responsible for ER-compartmentalization of eIF4E. mTORC1 activity reduction in growth retarded and mitochondria detethered cells influences the cell fate by acting on miRNA-mRNA axis. This is a unique way how mitochondrial activity is linked with protein translation and gene repression control in mammalian cells.

## Introduction

Micro RNAs (miRNAs) are small 20-24 nucleotide long non-coding RNAs that post-transcriptionally regulate the expression of their cognate mRNAs by base pairing and thereby regulate majority of genes in higher animals and plants. The miRNA repressed messages either undergo degradation or stored in specific cellular sites like P-bodies or stress granules [1–3]. Interestingly, importance of cellular structures and organelles in gene regulation has only emerged recently and these structures found to play an important role in controlling the activity of miRNAs. It has also been shown in current literature that miRNA repression and target RNA degradation are both temporally and spatially uncoupled processes [4]. Therefore, starting from biogenesis of a miRNA through its turnover requires involvement of various cellular organelles as they serve as destinations of miRNAs and respective target mRNAs under specific phases [5–8].

miRNAs bind to the 3’UTR of a target mRNA with pairing specificity [9] and downregulate the expression of proteins from target mRNAs by inducing deadenylation and decay of the same or by storing them in specialized RNA granules, the RNA processing bodies or P-bodies, in a translationally dormant state[10, 11]. miRNAs function in the form of ribonucleoprotein complexes-miRISCs (miRNA-induced silencing complexes). Components of miRISC and repressed mRNAs are also enriched in the P-bodies [12, 13]. P-bodies also interact with MVBs and targeting of miRNPs and repressed mRNAs to MVBs is perceived as a prerequisite for subsequent re-localization of miRNPs and repressed messages to P-bodies [4, 14]. In eukaryotes, degradation of mRNAs are usually initiated by shortening of the 3′ poly A tail, which leads to the removal of 5′ cap structure by a Dcp1/Dcp2 enzyme complex followed by 5′-3′ exonucleolytic digestion by XRN1 [15, 16]. The RNA decay machinery is reported to be accumulated in P-bodies [13, 17]. The concept of P-body involvement in mRNA storage function has been supported in a recent report where the authors have identified several miRNA-targeted mRNAs in biochemically purified P-bodies[18, 19].

Mitochondrial membrane potential not only helps production of ATP through oxidative phosphorylaton but it is also involved in miRNA-activity regulation. Interestingly, mitochondrial membrane potential disruption has implication in miRNA-mediated repression process [20] and it has been shown that mitochondrial membrane potential is required for Endoplasmic Reticulum (ER) and endosome-tethering of mitochondria that in turn controls miRNA activity in mammalian cells [6]. ER have been reported as sites of miRISC-mediated target RNA recognition and *de novo* miRNP formation [5, 21] while Multivesicular Bodies (MVBs) that interact with RNA processing bodies are the important sites for target RNA degradation [22]. Additionally, MVBs or late endosomes can merge with plasma membrane to export various cargoes through the formation of extracellular vesicles known as exosomes. These lipid bilayer sequestered bodies are small, 30-to 100-nm size vesicles considered as primary means of intercellular communication [23–26]. Multitude of miRNAs along with its associated proteins, such as Ago2 and GW182, are reported to be present in exosomes [27]. As of yet, the significance of miRNA export via exosomes and its regulation in addition to its role as a inter-cellular signalling factor for the maintenance of cellular homeostasis is only partly understood [28, 29].

Many alterations in cellular metabolic processes are greatly driven by changes in mitochondrial function and homeostasis. Classically, mitochondrial dysfunction has been implicated in cellular senescence mainly by promoting oxidative damage that induces cell-cycle arrest [30, 31]. However, emerging data suggests that other mitochondrial-dependent factors also play important role in the induction of senescent phenotypes [32].

mTOR (Mammalian target of rapamycin) plays a crucial role in controlling mitochondrial metabolism. Inhibition of mTOR activity leads to reduced mitochondrial respiration and enhanced aerobic glycolysis [33]. Another observation suggests mTORC1 stimulates mitochondrial biogenesis and activity and translation of mitochondria-related mRNAs, mediated by eIF-4E-BPs [34]. In reverse, how mitochondrial dysfunction influences mTORC1 activity to regulate cellular translational and miRNP machineries or their trafficking remains to be understood.

We report depolarization of mitochondria and its reduced dynamics, observed in growth arrested cells, are functionally coupled with retardation of miRNPs movement from polysomes to MVBs. This happens in conjunction with a uncoupling protein 2 (Ucp2) dependent mitochondrial depolarizationan and subsequent accumulation of miRNAs in polysome, that in turn cause the reduced exosomal miRNA export and P-body targeting of Ago proteins. Interestingly, reduced mTOR activity driven defective compartmentalization of translation initiation factor eIF4E to polysomes attached with the rER membrane causes the retarded translation of protein that in turn aborgates intracellular shuttling and subsequent export of miRNAs in mammalian cells.

## Results

### Reduced mitochondrial dynamics in growth retarded mammalian cells

Previously we have explored the status of miRNA machineries in cells that are growth retarded. These growth retarded High Density Culture (HDC) cells are morphologically and biochemically different from the proliferating Low Density Culture (LDC) cells [28]. To investigate the cause of the defective miRNA-metabolism happening in the HDC cells, we have explored the status of energy metabolism in both HDC and LDC cells to find its importance in defective miRNP machineries observed in HDC cells. It was important in the context of our previous observations where we have found that mitochondria play a key role in determining the activity of miRNPs in mammalian cells. In mammalian macrophage and also in non-macrophage cells mitochondrial detethering with endoplasmic reticulum has shown to cause increased miRNA stability that are also observed in HDC cell [6, 28].

To ascertain the effect of growth status on mitochondrial morphology and activity in human cells, HeLa and MDA-MB-231 cells were grown separately to LDC (25-40%) and HDC (100%) states. For assessing the mitochondrial status, the cells were transfected with a mitochondria targeting GFP (Mito-GFP) and an ER-targeting variant of DsRed2 (DsRed2-ER) and were studied microscopically. Evidently, a significant variation in shape and structure of mitochondria between HDC and LDC state cells were observed. In LDC condition, mitochondria appears as long filamentous structure whereas, in HDC condition they are predominantly punctate and spherical. However, ER looses most of its visible reticular structures in HDC state cells (Figure 1A; Supplementary Figure S1A). Moreover, when we quantified the length of individual mitochondrial filaments using 3D surface reconstructions of the confocal images taken for both LDC and HDC HeLa as well as MDA-MB-231 cells, we observed a mean length of ~6 μm in HDC condition as compared to ~20 μm long mitochondrial filaments in LDC cells (Figure 1A-B Supplementary Figure S1B). To connect the effect of this structural change of mitochondria to its dynamics in growth retarded cells, in the next phase of studies, we explored the spatial dynamics of mitochondria and ER organelles in both HDC and LDC cells by live cell imaging of the events. The spatial dynamic of mitochondrial structures of LDC cells was clearly different from that of HDC cells. While, mitochondria in LDC cells showed dynamic and frequent interaction with ER along with active mitochondrial fusion and fission in LDC cells, HDC mitochondria showed rather a sluggish dynamic and are of more punctate structures with negligible amount of mitochondrial fusion or ER overlaps (Figure 1C-D; Supplementary Figure S1C and Videos S1-4).

**Figure 1.**
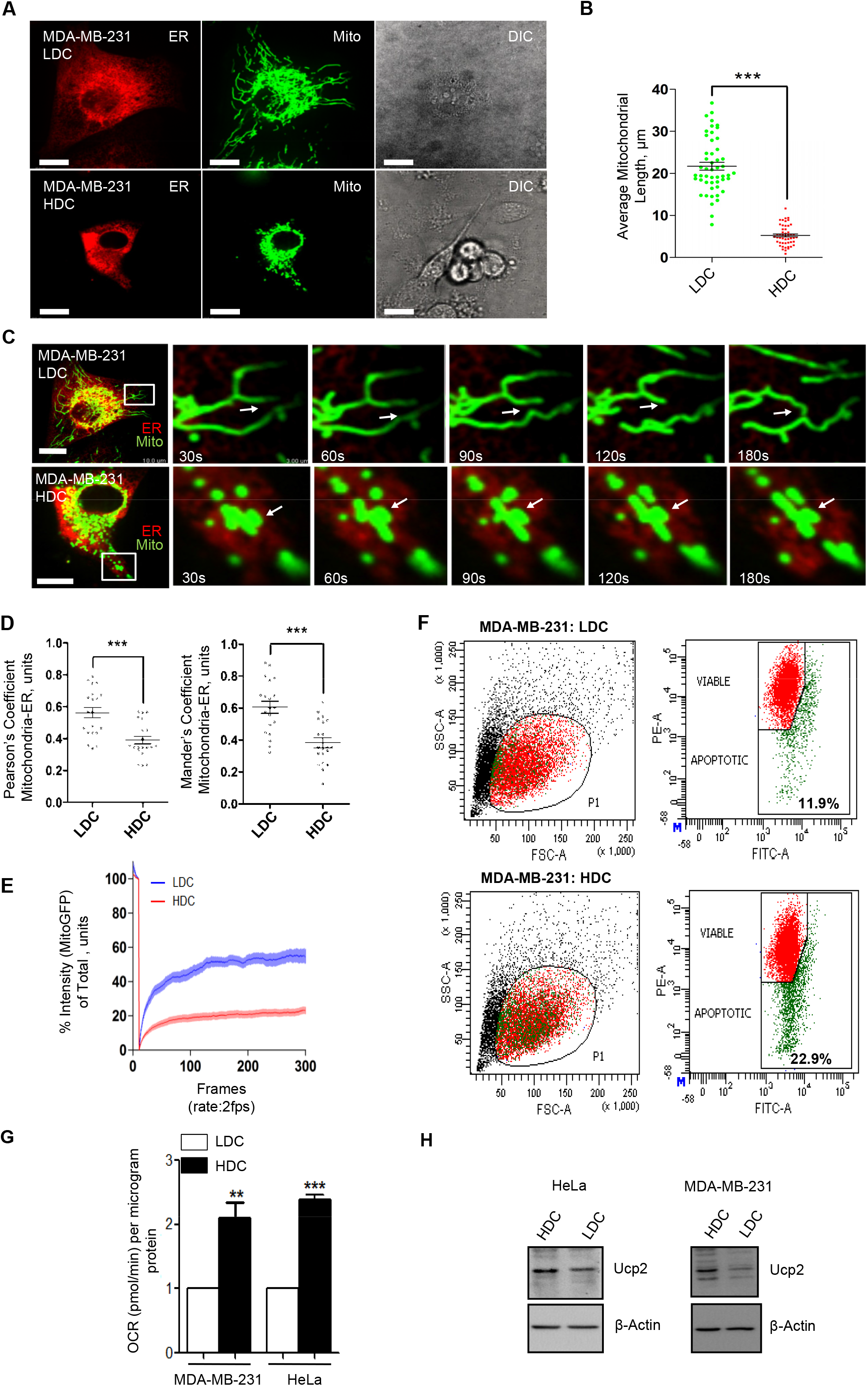
Alteration in mitochondrial morphology, dynamicity and membrane potential in growth retarded mammalian cells. **(A)** Representative pictures of MDA-MB-231 HDC and LDC state cells showing ER and mitochondrial structures. Cells expressing a mitochondrial targeting variant of GFP (MT-GFP, green) and an ER targeting variant of DsRed (pDsRed2-ER, red) were used. Scale bars are of 10μm length. (**B)** Mitochondrial size quantifications were done for LDC or HDC MDA-MB-231 cells. (**C)** Representative frames LDC or HDC MDA-MB-231 cells imaged live. Microscopy was done for a total of 3 mins at 1 fps. Cells expressing a mitochondrial targeting variant of GFP (MT-GFP, green) and an ER targeting variant of DsRed (pDsRed2-ER, red) were used. Bars measure 10μm. The ROI as depicted is 10X zoomed. Also see Video 1 and 2. (**D)** Pearson’s (*left*) and Mander’s (*right*) coefficients of mitochondria-ER colocalization in HDC and LDC cells. Calculation based on the images taken with cells described in panel **C**. (**E)** FRAP analysis of GFP-positive mitochondria in HDC or LDC MDA-MB-231 cells. Cells expressing a mitochondrial targeting variant of GFP (MT-GFP, green) were used. FRAP analysis from 100 cells (n>4) for indicated experimental sets. MT-GFP was photobleached and images were acquired at 2 frames per second for 300 frames. The mean intensity values of different regions of interest (ROI) thus obtained was fitted on a scale of 0 to 100. Mean intensity level of designated ROI before bleaching step taken as 100 and the value of Mean intensity in ROI just after photobleaching serving as 0. The values were plotted for a total of initial 300 frames. Shown are the mean and s.e.m for n>20.(**F)** Representative plots depicting flow cytometry based quantification of JC-1 stained HDC or LDC HeLa cells. Percent of cells in zone with low JC-1 Fluorescence (green) were measured and shown as depicted. (**G)** OCR (Oxygen Consumption Rate) determination in HDC and LDC HeLa cells. Quantification of OCR in HDC or LDC MDA-MB-231 and HeLa cells from 5 different experiments. (**H)** Representative western blot analysis of Ucp2 protein level in HDC or LDC HeLa or MDA-MB-231 cells. β-Actin served as loading control protein. For measuring statistical significance of the results, p values are calculated by Student’s t test, and one, two, and three asterisks represent p values less than 0.05, 0.01, and 0.001, respectively.

The structural alterations that we observed with the mitochondria of HDC or growth retarded cells could be due to a potential depreciation in inter-mitochondrial exchange happening specifically in HDC state cells. We performed FRAP based analysis to determine whether the mitochondrial content exchange in LDC and HDC cells are different between these two different status of the cell. This FRAP based analysis was utilized as a way to reconfirm the effects of cell densities on mitochondrial dynamics and we observed effects of cell growth status on mitochondrial fusion, mitochondrial length as well as mitochondrial exchange dynamics in human cells. Consistent with above mentioned observations, it was noted that the FRAP recovery rate of Mito-GFP in HDC cells was half that of in LDC cells, depicting a functional loss of mitochondrial dynamics and reaffirming the data we obtained for mitochondrial length shortening in HDC cells (Figure 1E).

### Reduced mitochondrial membrane potential in growth retarded cells

Mitochondrial membrane potential (ΔΨ_M_) modulation is integrally linked with changes in its structural and movement dynamics [35]. Therefore, it was not imperative to check the effect of cell density not only on structural and movement dynamic of mitochondria observed in HDC cells but also on the ΔΨ_M_, the mitochondrial membrane potential, in LDC and HDC cells. The assay based on binding of the dye JC-1 with polarized mitochondria was performed to measure the changes in ΔΨ_M_ due to increase in cellular density and a reduction in ΔΨ_M_ was observed in HDC cells compared to LDC state cells. The depreciation in ΔΨ_M_, measured by changes in uptake of JC-1, a cationic dye, was substantial (Figure 1F). The depression in ΔΨ_M_ was measured also by a more active and sensitive manner in Oxygen Consumption Rate (OCR) measurement, that reflects a sensitive and dynamic measure of the changes occurring in ΔΨ_M_. OCR based measurement of ΔΨ_M_, though indirect, is an excellent method of monitoring active variations in mitochondrial functional state. Distinct elevation in the levels of OCR in HDC condition cells of both HeLa as well as MDA-MB-231 was observed reflecting a lowered ΔΨ_M_ in HDC state cells (Figure 1G). Ucp2 is an endogenous mitochondrial uncoupler that is ubiquitously present across different tissues and we found increased expression of this protein in HDC state cells. As reported before, the increased expression of Ucp2 therefore could account for the lower ΔΨ_M_ observed in HDC state cells (Figure 1H).

### Mitochondria depolarization by FCCP causes accumulation of miRNAs and Ago2

It has shown previously that mitochondrial depolarization and detethering with ER, caused by the pathogen *Leishmania*, is associated with increased accumulation of miRNPs in host macrophages [6]. We have observed similar increase in miRNP content in HDC stage cells that also has depolarized mitochondria. Therefore, to establish a causal relationship between mitochondrial depolarization and miRNP accumulation, it was important to check the fate of the miRNPs upon induced depolarization of mitochondria. We used a specific chemical blocker of ΔΨ_M_, FCCP, to dissect the effect of ΔΨ_M_ reduction on miRNA activity and stability. FCCP treatment resulted in a robust loss of ΔΨ_M_ of treated cell mitochondria, as measured by JC-1 based flow cytometry assays (Figure 2A). Further on, when we used FCCP to depolarize mitochondria of HeLa cells, a large increase in the miRNA levels were observed as evidenced by the qRT-PCR based quantification of mature let-7a or miR-21 miRNAs in HeLa cells (Figure 2B, *left*). Increased level of an exogenously expressed liver specific miRNA-122 in HeLa cells was also noted with progressive FCCP treatment (Figure 2B, *right*). Furthermore, the *in vitro* target cleavage activity of miR-122 containing FLAG-HA-Ago2 miRISC purified from HeLa cells co-expressing miR-122 and FLAG-HA-Ago2 was measured. miRISC from FCCP treated cells showed elevated activity, that was consistent with the increased stabilization and Ago2 association of the miR-122 mimic, as observed in HDC state cells (Figure 2C-D). Evidently, previous studies have depicted the involvement of cellular organelles like rER in controlling miRNA life cycle [5, 21, 28, 36]. We checked the association of miRNPs with biochemically isolated rough ER fragments, the microsomes, in control and FCCP treated cells. Evident from the measurement of amount of Ago2 protein fractionated with the microsome, FCCP treatment increased the amount of Ago2 with microsomes (Figure 2E). To ascertain the more specific subcellular structures where the miRNP increase was occurring in FCCP treated cells, Optiprep^®^ based gradient was used for separation of cellular organelles [21]. Measurement of miRNA levels in individual fractions suggest elevated association of let-7a in the fractions, known to get enriched for endoplasmic reticulum [21], upon FCCP treatment as it was observed in HDC stage cells or in cells expressing Ucp2 exogenously[6, 28] [Supplementary Figure S2A-B].

**Figure 2.**
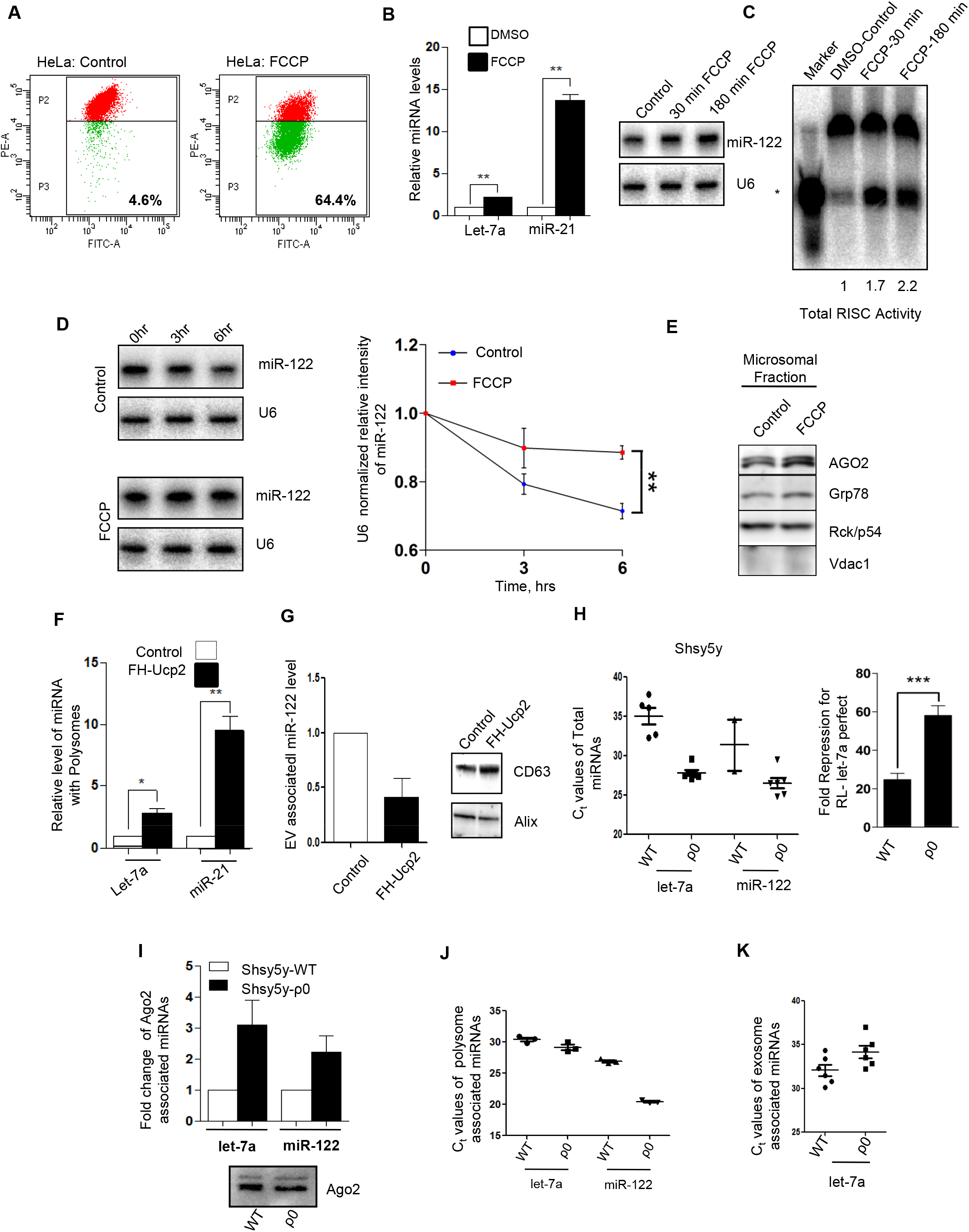
Mitochondrial depolarization causes the microsomal accumulation of miRNPs and reduced exosomal export of miRNAs. **(A)** Representative plots using flow cytometry based quantification of JC-1 stained control or FCCP treated HeLa cells. Percent of cells in zone of low JC-1 Fluorescence (green) were measured and shown as depicted. (**B)** This shows the levels of miR-21 and let-7a in FCCP treated cells. The values were estimated by qRT-PCR based method in control and treated cells. Relative levels of miR-122 were estimated by Northern blotting for HeLa cells treated with FCCP for indicated time periods (*right panel*). U6 was used as loading control for Northern blots and used for normalization in qRT-PCR based miRNA content quantifications. (**C)** Representative autoradiogram elucidating variation in RISC cleavage activity associated with Ago2 isolated from HeLa cells treated with FCCP for indicated time periods. *In vitro* RISC cleavage assay was done with isolated Ago2-miR-122 miRNPs from HeLa cells treated with FCCP for indicated time periods with FH-Ago2 expression construct and pmiR-122, a miR-122 expression plasmid. The activities were measured and quantified in an in vitro RISC cleavage reaction using 5’-γ^32^P labelled miR-122 target RNA as substrate. Values below the blot indicate relative RISC activity measured by estimating the cleaved band intensities against total amount of substrate used (uncleaved and cleaved RNA taken together) measured using densitometric analysis. Intensities were normalized to immunoprecipitated (IPed) FH-Ago2 levels. (*) indicates cleaved product. A 21 nucleotide long radiolabeled DNA oligo was used as size marker. The immunoprecipitated FH-Ago2 intensities have been used for calculating specific activity RISC cleavage and depicted by the respective quantified values. (**D)** Mature miR-122 decay rate estimated by Northern blotting (representative blot shown in the *left panel*) upon FCCP treatment for indicated time periods for HeLa cells. Band intensities from the northern blot for respective treatment times were normalized by U6 values and were used to plot the decay curve (*right panel*). Relative values were plotted for the respective time points. (**E)** Hela cells treated with FCCP or DMSO (Control) for a time period of 3 hours were used for the isolation of the total microsomal fraction by CaCl_2_ based separation from crude microsomal fraction. Representative images of western blot analysis for indicated proteins from FCCP treated microsomal fractions of HeLa cells. Absence of Vdac1 confirms the absence of mitochondrial contamination. (**F)** Polysomes from cell extract were extracted. The relative enrichment of the miR-21 and let-7a with polysomal fractions are plotted to determine the effect of Ucp2 expression on the polysomal microRNA level in mammalian cells. (**G)** qRT-PCR based relative quantification of exogenously expressed miR-122 in HeLa cells expressing FH-Ucp2. Values were normalized against respective miRNA levels in control cells. Shown are the mean and s.e.m from at least four independent experiments. Representative western blot analysis of exosomal marker CD-63 and Alix in the exosomal fraction of HeLa cells expressing FH-Ucp2 or control plasmid. Normalization was done against CD63 levels. (**H)** qRT-PCR based C_t_ values of endogenous let-7a or exogenously introduced miR-122 levels in WT or ρ0 SH-SY5Y cells. Shown are the mean and s.e.m from at least two independent experiments. In the right panel, *in vivo* let-7a miRNA activity in WT or ρ0 SH-SY5Y cells is shown. Relative repression level was measured by renilla luciferase luminescence in cells expressing luciferase based let-7a reporter with one let-7a perfect site, RL-Con, a reporter without let-7a site was used as control. **(I)** Immunoprecipitation of endogenous Ago2 was performed to estimate changes in Ago2-associated miRNA levels of WT or ρ0 SH-SY5Y cells. Quantification was done by qRT-PCR–based estimation. Values were normalized against amount of Ago2 immunoprecipitated and plotted against values of WT SH-SY5Y samples considered as unit. Representative blots showing immunoprecipitated endogenous Ago2 level used for normalization of respective miRNA levels. (**J-K)** qRT-PCR based C_t_ values of polysomal fraction (J), or exosomal fraction (K), associated endogenous let-7a or exogenously introduced miR-122 levels in WT or ρ0 SH-SY5Y cells. Shown are the mean and s.e.m from at least two independent experiments. p values are calculated by Student’s t test, and one, two, and three asterisks represent p values less than 0.05, 0.01, and 0.001, respectively. Shown are the mean and s.e.m for n>3.

### Reduced exosomal export of miRNAs in mitochondria depolarized cells

Increased retention of miRNAs with the polysomes, observed in HDC state cells, can be contributed by reduced mitochondrial activity and membrane potential. To test the fate of miRNAs in mitochondria depolarized cells, levels of miRNAs associated with polysomes was measured in HeLa cells expressing FH-Ucp2, a protein known to uncouple mitochondrial membrane potential[6]. There has been substantial increase in polysome attached miRNA content in FH-Ucp2 expressing LDC stage cells (Figure 2F). High retention of miRNA with polysomes that is associated with retarded export of miRNAs from HDC state cells and low levels of EV or exosome associated miRNAs in EVs isolated from HDC cell culture supernatant has been documented earlier [28]. Increased stability of miRNAs observed in HDC state human cells is primarily contributed by the retarded extracellular export of miRNAs happening in HDC state cells with increased miRNA retention with rER and polysomes. Like in HDC cells, expression of Ucp2 in LDC stage cells reduces the miRNA export via EVs (Figure 2G).

The reconfirmation of mitochondrial abnormality-driven defective extracellular miRNA export was strengthened in subsequent experiments where mitochondria defective ρ0 cells, an effective model to study neurodegerative disease, were used for measurement of miRNA activity and levels[37]. As expected we detected an increased cellular miRNA level in Shsy5y-ρ0 cells compared to the wild type cells (Figure 2H). An increased activity of miRISC observed was also consistent with increased cellular miRNA content (Figure 2H). The increased miRNAs found to be associated with Ago2 and polysomes (Figure 2I-J). It was accompanied by a reduced export of miRNA let-7a in ρ0 cells (Figure 2K). It has been observed earlier that mitochondrial detethering with ER can also cause increased miRNAs levels in mammalian cells [6]. We observed similar loss in mitochondria-ER interaction in ρ0 cells (Supplementary Figure S3A-B). This observation was consistent with the idea that the mitochondrial detethering caused by mitochondrial depolarization in mammalian cells is responsible for the effect that the depolarization has on the cellular and exosomal miRNA content in HDC cells. Indeed we also observed an expected decreased mitochondria-ER interaction in HDC cells (Figure 1D), which was consistent with the notion of mitochondrial depolarization induces loss of ER and mitochondria contacts[6].

### Mitochondrial depolarization affects P-body targeting of Ago2 in human cells

Previous reports clearly enunciate elevated miRNA stability in human cells upon Ucp2 upregulation and reduction in ΔΨ_M[6]_. Moreover, the effect of ΔΨ_M_ on Rck/p54 positive RNA processing body or P-bodies has been partly addressed in a previous study [38]. Interestingly, it has been observed that the miRNA-mediated clearance of target RNA get impaired in HDC cells where the target RNA is more stable than in proliferating cells[28]. This is consistent with a reduced trafficking of the repressed mRNAs to P-bodies, where they get degraded. In HDC cells polysome sequestration of both miRNA and its target messages can account for the increased mRNA stability found in HDC or in mitochondria depolarized cells. To discover the link between loss of ΔΨ_M_ and P-body status, microscopic analysis were done in cells treated with either FCCP or exogenously expressed FLAG and HA tagged version of Ucp2 (FH-Ucp2)-upregulation of which cause disruption of mitochondrial membrane potential [6]. A distinct loss in visible Ago2 positive P-bodies was observed in cells either treated with FCCP or expressing FH-Ucp2 (Figure 3A-B). As expected we also documented a reduced number of P-bodies in SH-SY5Y ρ0 cells (Supplementary Figure S4C).

**Figure 3.**
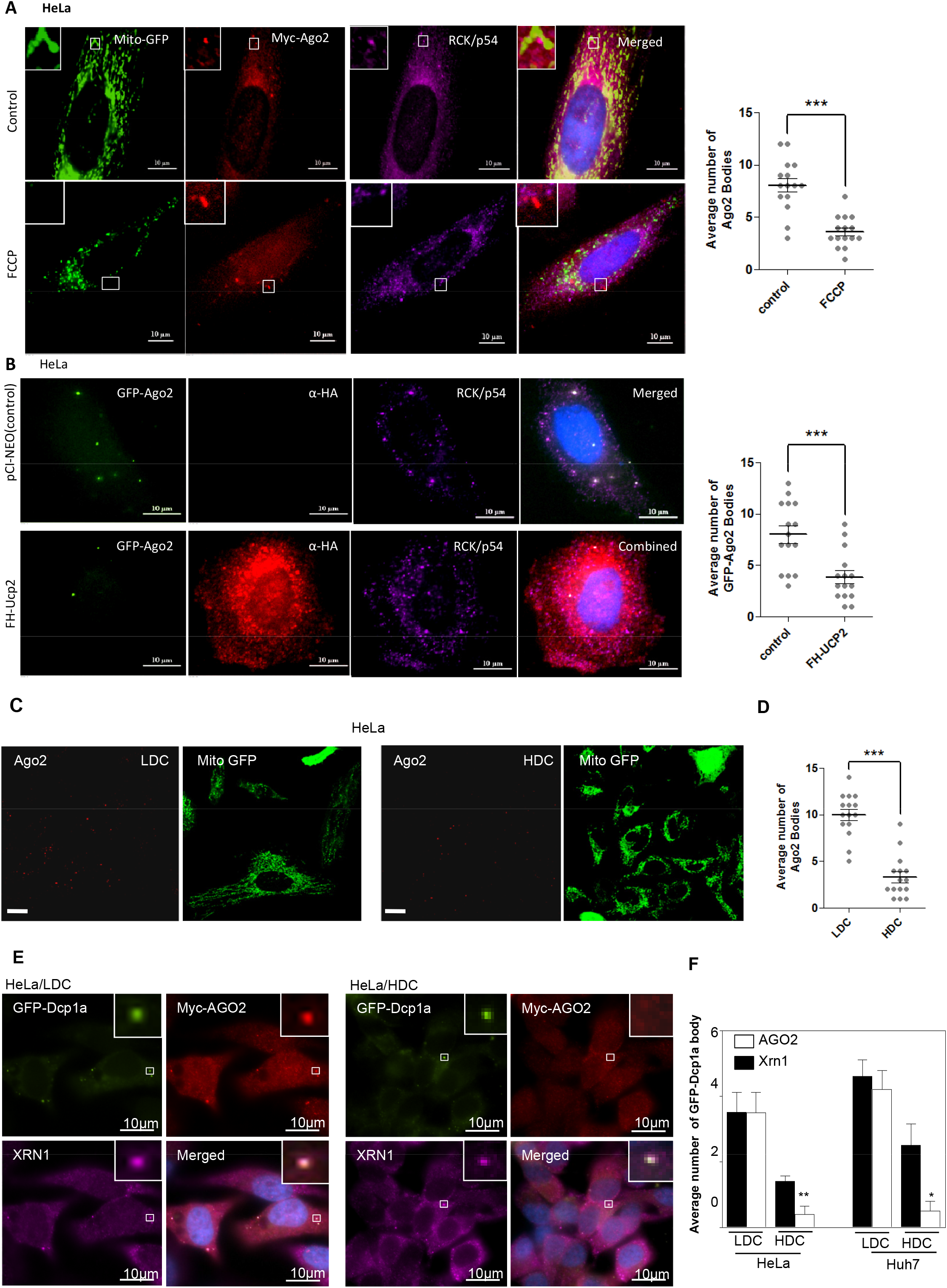
Effect of mitochondrial depolarization on P-bodies. **(A)** FCCP treatment affects P-bodies. Representative frames of mitochondria in HeLa cells treated with FCCP. Cells expressing a mitochondrial targeting variant of GFP (MT-GFP, *green*) and a myc-tagged variant of Ago2 were used. Ago2 were labelled using anti-myc (*red*) and antibodies against endogenous Rck/p54 (*Magenta*) was used to label them by indirect immunofluorescence. DAPI was used for depicting nucleus. Bars measure 10μm. (*left*) Quantification of myc-Ago2 positive bodies normalized to the number of cells (*right*). (**B)** Effect of Ucp2 expression on P-body status in HeLa cells. Representative frames of Ago2 in HeLa cells expressing FH-Ucp2. Cells expressing an Ago2 variant of GFP (Ago2-GFP, *green*) and FH-Ucp2 (*red*) were used. HA-Ucp2 were labelled using anti-HA and antibodies against endogenous Rck/p54 (*Magenta*) was used to label P-bodies by indirect immunofluorescence. DAPI was used for depicting nucleus. Bars measure 10μm (*left*). Quantification of GFP-Ago2 bodies normalized to the number of cells *(right)*. (**C)** Representative confocal microscopy frames depicting mitochondria in HDC or LDC HeLa cells. Cells expressing a mitochondrial targeting variant of GFP (MT-GFP, green) were used. Endogenous Ago2 was labelled with Alexa-594. Bars measure 5μm. (**D)** Estimation of Ago2 bodies was done from frames as depicted in *panel* ***C*** and normalized with number of cells using Imaris. (**E)** Representative fluorescence microscopy frames of P-body markers in HDC or LDC HeLa cells. Cells expressing a GFP tagged variant of Dcp1a (*green*) (GFP-Dcp1a) and a myc tagged variant of Ago2 (Myc-Ago2) were used. Anti-Myc antibody was used to label myc-Ago2 and it was labelled with Alexa 594 (*red*). Endogenous XRN1 was labelled with Alexa-647 (*violet*). DAPI staining was used to depict nuclei. Bars measure 10μm. (**F)** Estimation of Dcp1a bodies colocalizing with protein factors as depicted in HDC or LDC conditions of HeLa cell from frames, as depicted in a representative picture in *panel **C*** and normalized against number of cells. p values are calculated by Student’s t test, and one, two, and three asterisks represent p values less than 0.05, 0.01, and 0.001, respectively. Shown are the mean and s.e.m from at least five independent experiments.

### Reduced P-body dynamics and Ago2 localization also happens in growth retarded cells

Does impaired intracellular trafficking of miRNAs also resulted in defective P-body localization in HDC stage cells? To answer, the effect of cell density on P-bodies was studied in the context of HDC or LDC stages. There was a profound decrease in the number of visible Ago2 granules (Figure 3C-D). To characterize further the effects of cellular confluency on the status of P-bodies, we conducted immunofluorescence analysis of two established P-body components, Dcp1a and XRN1. In other cases we used tagged versions of Dcp1a or Ago2 to track and document P-body status. The decapping (Dcp) proteins are conserved P-body components in yeast, *C. elegans*, *Drosophila* and mammalian cells [39]. Dcp2 is the decapping enzyme that removes the 5’ cap from the mRNA destined for degradation. Dcp1a is a stimulator of Dcp2 activity [39]. XRN1 gene encodes a member of the 5’-3’ exonuclease family and is a well conserved marker of P-body structures [40, 41]. To further probe the loss of distinct P-bodies with cell confluency, HeLa cells were transfected with Myc-Ago2 and GFP-Dcp1a, further on these cells were co-immunostained with XRN1 to monitor the localization of Ago2 in such conditions. In LDC HeLa cells most of the GFP-Dcp1a bodies were positive for Myc-Ago2 while in HDC cells Ago2 co-localization with GFP-Dcp1a was visibly impaired. Additionally, in HDC HeLa cells the average number of discrete P-bodies was low (Figure 3E-F).

P-body components are known to exchange their content with the cytosol and most of the P-body markers are also ubiquitously found in the cytoplasm [42]. To ascertain whether the exchange dynamics of cargo is modulated in cells grown to confluent states, we measured fluorescence recovery after photobleaching (FRAP) of GFP-tagged P-body protein markers in cells expressing GFP-Dcp1a or GFP-Ago2. Majority of P-bodies in HDC and LDC HeLa cells were found to be dynamically distinct though similar recovery for GFP-Dcp1a to P-bodies were seen in both HDC and LDC cells. In contrast, recovery of GFP-Ago2 to P-bodies was reduced in HDC cells compared to LDC cells (Supplementary Figure S4A). The linear velocity and radii of GFP-Ago2 positive P-bodies were distinctly higher in LDC than HDC HeLa cells (Supplementary Figure S4B-C).

### Restoration of mitochondrial potential rescues intracellular miRNP shuttling and P-bodies

The ΔΨ_M_ mediated effects upon P-body number, Ago2-localisation to P-bodies, polysomal sequestration of miRNAs and lowered miRNA export through EVs all occurs concurrently with a marked increase in Ucp2 protein levels. Moreover, many of these results are replicated in cells upon loss of ΔΨ_M_ through chemical blocking of mitochondrial membrane potential depicting a similar situation with exogenous expression of Ucp2 in mammalian cells. Therefore to specifically link Ucp2 upregulation with these effects on miRNP machineries and P-bodies observed in HDC cells, we used Genipin, which is a specific blocker of Ucp2 in human cells when Ucp2 level was already elevated in HDC stage cells. Successful restoration of ΔΨ_M_ in HeLa HDC cells was observed post Genipin treatment (Supplementary Figure S5A). Moreover, the effect on Ago2-positive P-body number as well as Ago2 co-localization with Rck/p54 was partially restored upon Genipin treatment of HDC cells (Supplementary Figure S5B-C). This rescue in P-body number happened due to a near complete reversal of Ucp2 protein levels in Genipin treated HDC cells (Supplementary Figure S5D). Furthermore, both polysomal enrichment of miRNA as well as lowered release of exosomal miRNAs from HDC cells were rescued to near LDC levels upon specific blocking of Ucp2 (Supplementary Figure S5E-F). Hence, mitochondrial depolarization is directly linked with miRNA function as well as its stability and an elevation of Ucp2 protein level and function occurs in growth retarded cells and can be restored by Genipin to restore miRNA levels and P-bodies.

### Mitochondrial detethering induced retention of miRNPs on rER is caused by reduced rER targetting of eIF4E

Previous studies have shown that ΔΨ_M_ regulates the mitochondria-ER tethering through Mitofusin2 (Mfn2) [43]. Moreover, loss of mitochondrial fusion in mammalian cells is associated with the punctate form of mitochondrial due to a resultant increase in mitochondrial fission [44–46]. Hence, we measured the cellular and polysomal association of miRNA and Ago2 protein in mouse embryonic fibroblasts either expressing the wild-type Mfn2 protein or without the Mfn2 encoding genes, to connect mitochondrial depolarization and detethering observed in HDC cells with reduced EV-mediated miRNA export. Interestingly, in the Mfn2 knock-out cells we observed a substantial increase in polysome associated miRNA and Ago2 protein levels with concomitant decrease in exosomal miRNA content in EVs isolated from those cells (Figure 4A-C).

**Figure 4.**
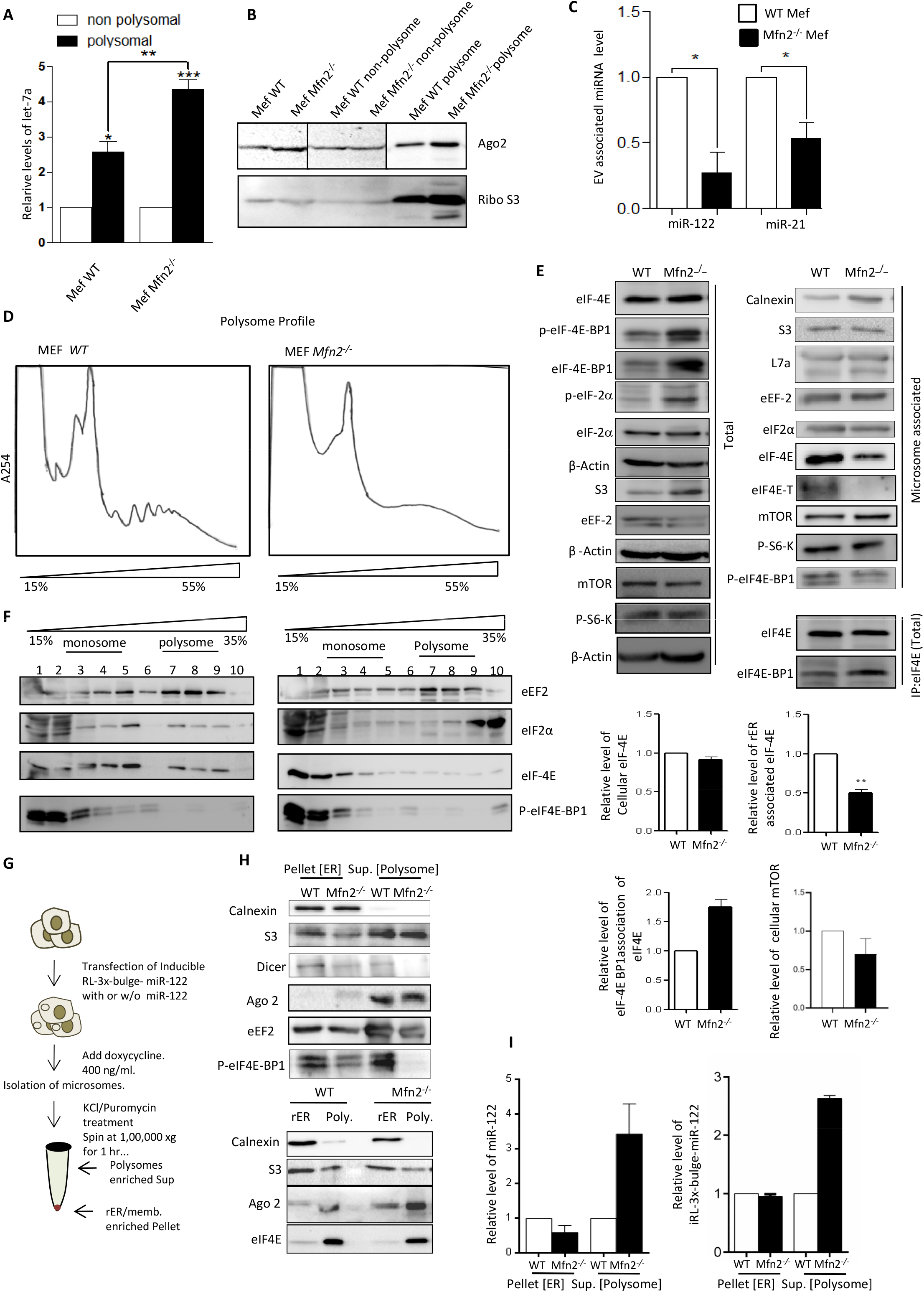
Defective targeting of eIF4E to rER attached polysomes to cause retarded intracellular miRNA trafficking and export. **(A** qRT-PCR based relative quantification of endogenous let-7a levels in polysomal and non-polysomal fractions obtained from one-step polysome separation done with MEFs of indicated genotypes. Values were normalized against U6 RNA and relative miRNA levels are shown. Values are the mean and s.e.m from at least four independent experiments. (**B**) Representative western blot analysis of Ago2 in total extract or polysomal and non-polysomal fractions after one-step polysome separation done with MEFs of indicated genotypes. For comparison between cells of different genotypes equal proteins were loaded while for comparison between subcellular fractions cell equivalent amount of each fraction were analyzed. Ribosomal S3 marks the presence of ribosomes which is enriched in the polysomal fraction. (**C**) qRT-PCR based relative quantification of exogenously expressed miR-122 or endogenous miR-21 levels in Extracellular Vesicles (EVs) derived from MEFs of indicated genotypes. Individual values were normalized against protein content of the EVs and relative values between cell types are shown. Values are the mean and s.e.m from at least four independent experiments. (**D**) Cell extracts from WT and Mfn2^−/−^ MEFs were isolated and analyzed on 15-55% sucrose density gradient. Gradient fractions were further collected and absorbance was monitored at 254nm and the respective absorbance profile graphs were plotted. (**E**) Relative abundance of different translation factors (eIF4E, P-eIF-4EBP1, P-eIF2α, eEF2 and eIF2α), mTOR or ribosomal protein S3 in cell extracts and microsomes derived from MEF WT and MEF Mfn2^−/−^ cells. Total or microsomal fractions were analyzed by western blotting. β-Actin and Calnexin were served as an internal control for cellular and microsomal samples respectively. Endogenous cellular eIF4E was immunoprecipitated and association of eIF-4EBP1 with immunoprecipitated materials was analyzed by western blotting in WT and Mfn2^−/−^ MEF extracts. Relative quantification of eIF4E in individual fractions or its interaction with eIF4E-BP1 was quantified by densitometric analysis of western blot data from multiple experiments. Relative mTOR levels with microsome were also quantified. (**F)** Extracts of MEFs (WT and Mfn2^−/−^) were isolated and analyzed on 15-35% sucrose density gradient and different fractions were collected and proteins were isolated. Distribution profile of different translation factors (viz. eEF2, eIF2α, eIF4E, p-eIF-4EBP1) were analyzed by western blots. (**G)** Schematic representation of KCl/Puromycin based extraction method for isolation of polysomes attached with the microsomal membrane from MEFs. (**H-I)** Microsomes from WT or Mfn2^−/−^ MEFs were treated with KCl/Puromycin to extract the translating polysomes. Further, proteins and RNAs were isolated from residual ER and solubilised polysome enriched fractions. Samples were analyzed by western blotting done to check distribution of miRNP proteins (Ago2 and Dicer) and translation factors after the extraction. Respective distribution of the ER and ribosomal marker proteins (Calnexin and S3) were also checked and was followed to ensure proper fractionation in panel **H**. In panel **I** Amount of miRNAs and mRNAs were also quantified by qRT-PCR based quantification against U6 or ribosomal RNA content. p values are calculated by Student’s t-test, and one, two, and three asterisks represent p values less than 0.05, 0.01, and 0.001, respectively. Shown are the mean and s.e.m for n>3.

What could cause the increased association of miRNPs with rER in Mfn2 negative cells? We observed a reduced amount of polysome in Mfn2^−/−^ cells (Figure 4D). We checked the distribution of several translation initiation regulatory factors along with the elongation factor eEF2, and detected a specific loss of eIF4E, the very important cap binding protein essential for translational initiation, on rER associated fraction (Figure 4E) [47]. Although in the cellular levels of eIF4E remain unchanged in both in wild type and Mfn2^−/−^ cells (Figure 4E). Interestingly, in Mfn2^−/−^ cells, eIF4E also showed an increased interaction with eIF4E-BP1 that may be due to an increased expression and decreased phosphorylation of eIF4E-BP1 ultimately causing an increase in its eIF4E binding in Mfn2^−/−^ cells (Figure 4E). The reduced recruitment of eIF4E to ER is consistent with poor translation observed in Mfn2^−/−^ cells that was revealed in the 15-35% sucrose density gradient analysis of cell extract for the components of protein translation machinery. Association of eIF4E with the heavier polysomal fraction was noted to be severally affected in Mfn2^−/−^ cells compared to wild type cells (Figure 4F). KCl-Puromycin treatment can extract attached polysomes from the ER membrane [5]. In subsequent experiments we have found association of most of the translation initiation factors including eIF4E with KCl-Puromycin soluble translating polysome fraction attached with MEFs derived microsomes (Figure 4G-H). This is consistent with the previous observations that suggest microsome attached membrane as the major site for translation regulation where the target message and miRNAs accumulates, interact and mRNAs either get translated or repressed. It has been shown previously that the low translatability of the mRNAs in HDC cells causes retention of the miRNPs with the rER attached polysomes to cause increased miRNA stability as effective translation is pre-requisite for miRNP recycling [28]. We also observed increased accumulation of target message and miRNA with the rER attached polysomes in Mfn2 negative cells (Figure 4I). Therefore mitochondial detethering of rER, that also observed in HDC cells, may directly cause lowering of miRNA turnover by targeting the initiation phase of translation caused by poor shuttling of cap binding protein eIF4E from cytoplasmic pool to the rER associated domain, a process controlled by its strong interaction with eIF4E-BP1 observed in mitochondria detethered cells. Interestingly the eIF4E-T(eIF4E-transporter), nuclear importer of eIF4E and the competitor of eIF4E-BP1 for eIF-4E binding, also found predominantly absent in Mfn2^−/−^ cells (Figure 4E)[48]. The data suggests eIF-4EBP1 might get increased access to inactivate eIF4E on this eIF4E-T–absent scenario in Mfn2^−/−^ cells.

We wanted to study the upstream regulatory kinase of eIF4E-BP1, mTOR also to understand the regulatory cascade. Here we could observe decreased mTOR expression on cellular as well as on microsomal levels in Mfn2^−/−^ cells (Figure 4E). Inhibition on mTORC1 complex activity leads to phosphorylation of eIF2α, which ultimately results in its inactivation and downregulation of global protein synthesis inside the cells [49]. Here we could also observe increased phosphorylation of eIF2α, correlating with defective translation status in Mfn2^−/−^ cells (Figure 4E).

### Reduced mTORC1 activity to restrict the relocalization of eIF4E to rER

To prove the importance of eIF4E relocalization to polysomes in controlling miRNA stability in mammalian cells, we had treated MDAM-MB-231 cells with Rapamycin, the mTORC1 inhibitor [47], and had noted the reduction mTORC1 phosphorylation and downstream activity evident in reduced in S6K phosphorylation in Rapamycin treated cells (Figure 5A). This was consistent with a reduction in the level of eIF4E-BP1 phsophorylation and relocalization of eIF4E to rER fraction (Figure 5A). With a reduction in eIF4E location to rER or microsomes, we documented an increase in miRNA content both in total and polysome associated pools upon Rapamycin treatment (Figure 5B-C). Like in HDC or HA-Ucp2 expressing cells, Rapamycin treatment also resulted in lower number of P-bodies in treated cells (Figure 5D)

**Figure 5.**
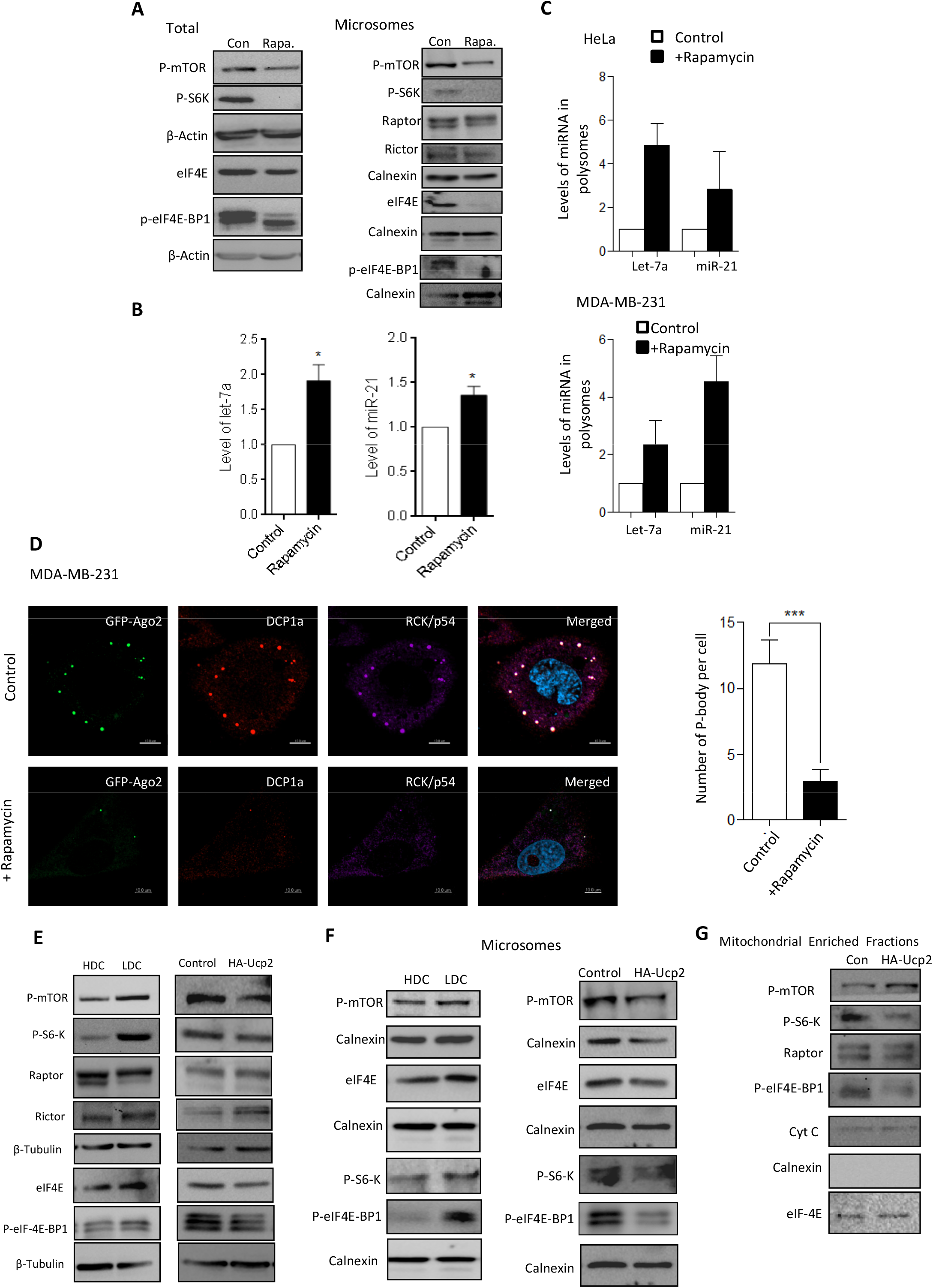
Rapamycin, the mTORC1 inhibitor, retarded eIF4E targeting to microsome and reduces miRNA turnover. Cellular and microsomal distribution of different translation factors (eIF4E, P-eIF4E-BP1, p-S6-K) and mTOR complex proteins (P-mTOR, Raptor and Rictor) after 16 hrs of 100 nM Rapamycin treated and Control (DMSO treated) MDAM-MB-231 cells. Proteins samples from total or microsomal fractions were analyzed by western blotting. Beta Actin or Calnexin were used as loading control for total or microsomal fractions respectively. (**B)** Quantitative RT-PCR data to check cellular let-7a and miR-21 levels after 16 hrs of 100 nM Rapamycin treatment compared to control (DMSO-treated) MDAM-MB-231 cells. U6 snRNA served as an internal control. (**C)** Quantitative RT-PCR data shows let-7a and miR-21 levels on polysomal fractions after 16 hrs of 100 nM Rapamycin treatment compared to control (no Rapamycin) of MDAM-MB-231 cells. U6 snRNA was used for normaliztion. (**D)** Representative confocal microscopy images showing P-bodies in untreated control or 100 nM Rapamycin-treated MDA-MB-231 cells. Cells expressing an Ago2 variant of GFP (Ago2-GFP, *green*) were used for treatment. Antibodies against endogenous Rck/p54 (*magenta*) and Dcp1a (*red*) was used to label P-bodies by indirect immunofluorescence. Bars measure 10μM. DAPI was used for depicting the nuclei. Quantification of number of P-bodies, normalized to the number of cells was done for untreated control or Rapamycin treated MDA-MB-231 cells as shown in the representative image obtained. (**E)** Expression of mTORC1, eIF4E, phoshorylated S6K and eIF4E-BP1 on LDC and HDC MDAM-MB-231 cells. β-tubulin was used for normalization. (**F)** Levels of translational initiation factor eIF4E and phosphorylated mTOR with microsome isolated from HDC or LDC cells and also in HA-Ucp2 cells compared to vector control transfected cells. Amount of microsomes used for analysis were evident from the calnexin levels in individual fraction. (**G)** Levels of mTOR and associated factors with mitochondria isolated from control and HA-Ucp2 expressing cells. Paired two-tailed Student’s t tests were used for all comparisons. P < 0.05 (*); p < 0.01 (**); p < 0.001(***). In (**B-C**) values are means from at least three biological replicates ± SD.

Mitochondrial dysfunction has been reported previously to cause rapid FRAP(mTOR)-dependent p70S6K deactivation [50]. In this report, it has been shown how FKBP12 the rapamycin-associated protein got associated with mitochondria to senses osmotic stress via mitochondrial dysfunction to control mTOR activity. In another report, low mitochondrial respiration has been found to be correlated with attenuation of mTOR pathway with decreased expression of p70S6K in triple-negative breast cancer cell lines [51]. We have done experiments in HDC and LDC cells to score the levels of phospho mTOR and eIF4E and noted a decrease on phospho mTOR content along with eIF-4E levels in HDC cells (Figure 5E). With isolated microsome or rER we observed a reduction of eIF4E content in HDC cells microsomes along with a reduced phospho mTOR levels (Figure 5F). Similar lowering of eIF4E was also noted in cells expressing FH-Ucp2 (Figure 5F). Therefore reduced mTORC1 activity is responsible for reduced phosphorylation of eIF4E-BP1 and free eIF4E required for miRNA trafficking and export in HDC cells. Similar decrease of microsome associated mTOR was also noted in cells depleted for Mfn2 (Figure 4E). The defective mitochondrial function is possibly responsible for the defective mTOR activation and reduced eIF-4E association with ER attached polysome-a point further supported in analysis done with mitochondrial fraction. Mitochondria associated mTOR was found to be increased in amount in HA-Ucp2 expressing cells with a decrease in Raptor amount. This can explain why the mTOR is less active in HA-Ucp2 expressing cells with a reduced P-eIF4E-BP1 and S6K phosphorylation and less eIF-4E association with microsomes (Figure 5G and Figure 6).

**Figure 6.**
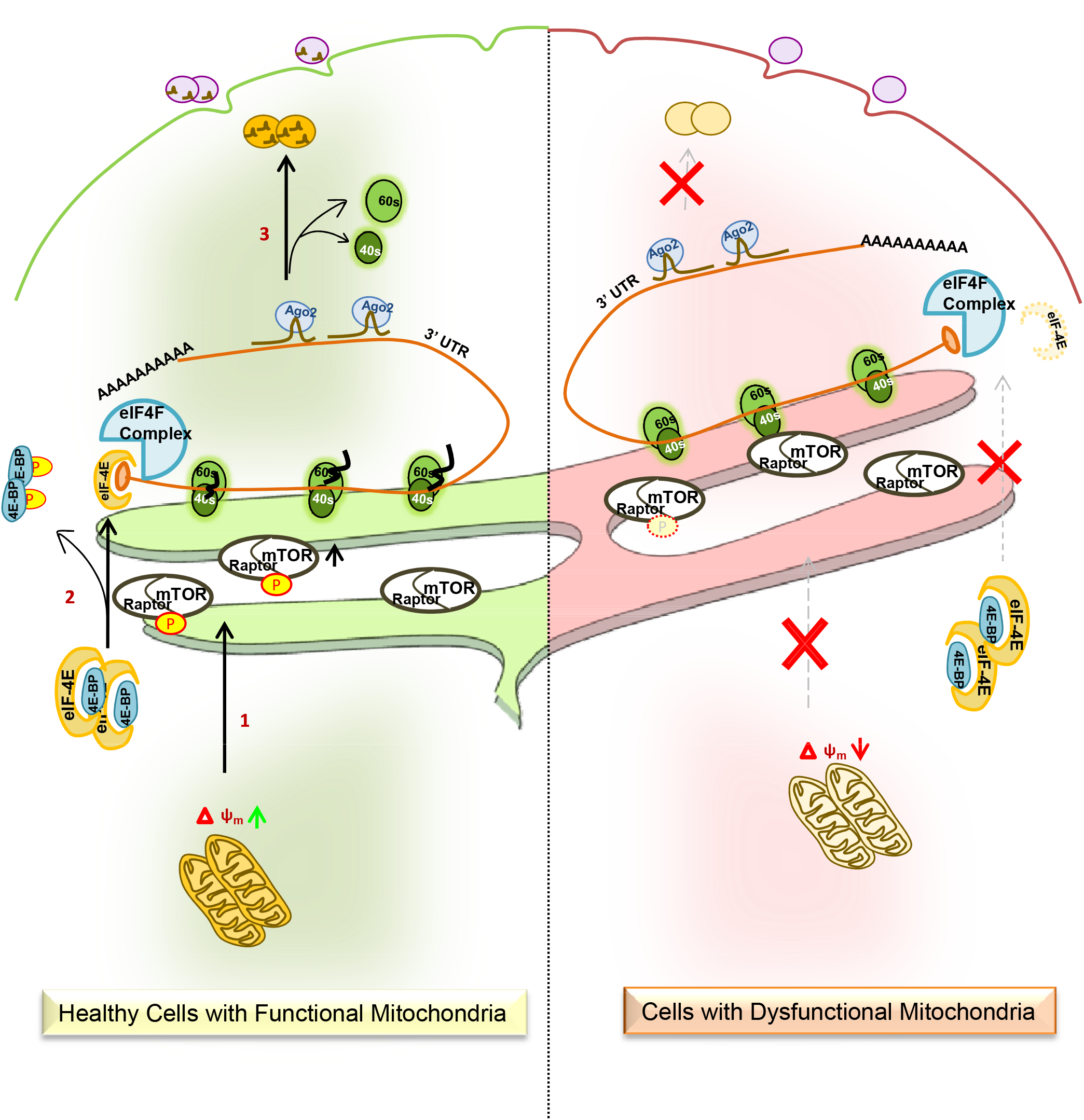
A schematic representation of how mitochondria controlled differential activation of mTOR governs intracellular trafficking and extracellular export of miRNAs in mammalian cells. Human cells with functional mitochondria shows proper activation of mTOR that leads to subsequent migration and phosphorylation of eIF4E-BPs on polysomes attached with rER to dissociate inactive eIF4Es and recruitment onto the active eIF4F translation initiation complex. This follows proper intracellular shuttling and extracellular export of miRNAs and dissociation of ribosomal subunits for another set of new cycle. In contrary to this, cells containing defective mitochondria (depolarized or detethered with rER) shows reduced activity of mTOR, that manifests in decreased migration and phosphorytaion of eIF4E-BPs with reduced active eIF4E level. Ultimately this causes enhanced ribosomal retention of miRNAs (28) and target mRNAs with poor intracellular trafficking on rER attached polysomes.

## Discussion

The incorporation of eIF4E into the eIF4F translation initiantion complex is reciprocally controlled through a family of inhibitor proteins, called the eIF4E-binding proteins (4E-BPs). eIF-4E mediates the attachment of eIF4F with 5’-cap structure of the mRNA to initiate cap-dependent translation in the cytoplasm. Phosphorylation of mTOR on rER membrane probably triggers a cue for eIF4E-BP1 to bring itself close to rER for phosphorylation. This in turn brings attached inactive eIF4E to rER membrane more effectively, after phosphorylation of eIF4E-BP1 by P-mTOR it gets dissociated with eIF4E and makes it active. Assembling of fuctional eIF4E on pre-initiation complex leads to proper functioning of translational & miRNP machineries could take place on polysomes attached with rER. Whereas in cells with defective mitochondrial condition or treated with an inhibitor, mTOR could not get properly activated that resulted in decreased rER migration of eIF-4E, which leads to reduced translation and increased stability & poor intracellular shuttling of mRNAs and the cognate miRNAs. Here we speculate eIF4E-BP1 senses the status of activated mTOR which ultimately determine the localization of eIF4E to regulate mRNA stability and miRNP turnover by influencing translation and miRNP trafficking. MiRNA activities are regulated through various different types of mechanisms and these regulations are important for normal cellular function. Amongst these, spatial sequestration of miRNA through the involvement of subcellular organelles like Golgi, ER and endosomes is found to be crucial [4, 7]. Though mitochondria has only been amongst the fringe players in this process while recent reports define mitochondria along with ER as the key players in this context [4, 5, 36]. Mitochondrial control of miRNA function and stability are primarily controlled by Ucp2 and Mfn2. Ucp2 alters the ΔΨ_M_ and affects tethering between mitochondria and ER that in turn found to be essential for the association and interaction between ER and endosomes. This association is evidently important for the transfer of target RNA loaded miRISC from ER, site of miRISC-target RNA interaction, to endosomes and MVBs where miRISC uncoupling and target RNA degradation occurs [4, 6]. Elevation of Ucp2 protein levels cause increased uncoupling of mitochondria which results in a cascade of events including defective interaction between these membranes. Moreover, polysomal miRISC interaction is known to be dependent on the translational speed and efficiency [28]. Thus mitochondria might be the key player in protein translation as potential supplier of localized ATP and hence, act as a determinant even during the earlier steps in miRISC nucleation process by modulating global translational rate. It would be interesting to estimate dependency of miRNA based repression process on the amount of mitochondrial ATP.

The importance of mRNA translatability is another important factor dominating the extracellular miRNA export and its stability. The increased stabilities of both miRNAs and its target messages were contributed by their retention with translationally "inactive" polysomes attached with rER membrane in HDC cells [28]. This could get reversed in proliferating cells. The importance of m7G cap-binding of a mRNA by eIF4E and its recruitment to mRNAs is prerequisite for continuous production of proteins from respective mRNAs. In previous papers, eIF4E-recruitment has been shown as one of the highly regulated steps in eukaryotic translation control where several competitor molecules including several cap binding proteins like eIF4E-T could inhibit the eIF4E interaction with 5’ Cap structure [52]. Recruitment of some of these inhibitory factors is even regulated indirectly by miRNPs bound to cis-acting elements on the 3’UTR of the same mRNAs [52].

Despite compelling evidence about the role of P-Bodies in the miRNA mediated mRNA surveillance; several issues still remain elusive in defining the miRNA-mRNA-P body axis. The role of the P-bodies as the degradation sites of mRNA and the mechanisms relevant to a particular context is an essential, yet difficult to answer question because the underlying networks are large, complex and only partially understood. Recently, discovery of miRNA-targeted messages in biochemically purified P-Bodies suggest rather a storage function of P-bodies that was originally predicted and supported in specific context. Since the disassembly of large P-bodies during high density of growth does not completely abrogate mRNA decay [28]; it remains to be explored if P-body function can be sufficiently backed up by microscopically invisible small-sized PBs (that may be unaltered in HDC state cells). Inhibition of miRNA production reduces P-body numbers in mammalian cells [53]. However, deployment of mRNAs from the translation machineries by miRNA-repressive action is considered a prerequisite for P-body localization of miRNP-targeted mRNAs. Maybe this process of spatial aggregation reduces the energetic burden on the cell by lowering the overall entropy of the system. Our study reveals the interesting changes in cellular P-body numbers and dynamics in response to environmental cues originating from cell-cell contact modulated by the cellular energetic landscape that primarily governed by mitochondrial functionality and status.

The ER-endosome association, an interaction regulated through Ucp2 mediated perturbation of mitochondrial membrane potential, is defective in growth retarded cells due to an elevation in Ucp2 protein levels. This, resulting in a loss of association between the pools of miRNPs present on ER and endosomes in growth retarded cells causes a subsequent rise in polysomal miRNA content with reduced P-Body targeting of miRNP components like Ago protein. Moreover, a concurrent decrease in miRNA content of extracellular vesicles was observed seen in both mitochondria compromised and HDC cells. Interestingly, this aberration occurs along with a marginal but significant rise in total exosomal membrane content. Thus, loading of the miRNA is dependent on the release of miRNPs from ER post its interaction with target RNAs. Moreover this type of block also affects the formation of visible P-bodies. Furthermore, Ucp2 driven alteration is sufficient for organelle interaction mediated miRNA regulation and therefore Ucp2 might potentially act as a key regulator of various cell fate determinant processes.

Overall it seems that mitochondria by regulating the organeller dynamics and inter-organeller exchange of materials plays a key role in controlling miRNP redistribution and activity modulation and are affected in situation where mitochondrial activities are heavily regulated by both intrinsic and extrinsic factors such as availability of nutrients and growth factors.

## Materials and Methods

### Cell Culture and treatment

Human HeLa, MDA-MB-231, MEFs, SH-SY5Y and Huh7 cells were grown in Dulbecco’s modified Eagle’s medium (DMEM) with 2 mM L-glutamine and supplemented with 10% heat-inactivated fetal calf serum (FCS). For all experiments cells were grown to 25-40% confluency states (LDC) or visibly full confluent (HDC) states unless specified otherwise [28]. Transfections were performed using Lipofectamine 2000 (Invitrogen) following manufacturer’s protocol. FCCP (500nM) treatment was done for indicated time points. Genipin and Rapamycin were used at 100 μM and 100nM concentration respectively[6].

### Immunofluorescence

For immunofluorescence, cells grown on 6 well tissue culture plate were transfected with 250 ng of GFP-Ago2, GFP-Dcp1a or N-HA-GW182 encoding plasmids[53]. The cells were split after 24h of transfection and subjected to specific experimental conditions. For immunofluorescence analysis, cells were fixed with 4% paraformaldehyde for 30 min, permeabilized and blocked with PBS containing 1% BSA and 0.1% Triton X-100 and 10% goat serum (GIBCO) for 30 min. After incubation with primary antibodies in the same buffer at desired dilution (see Information Table 1) overnight at 4°C and subsequent washing steps, secondary anti-rabbit or anti-mouse antibodies labeled either with Alexa Fluor® 488 dye (green), Alexa Fluor® 594 dye (red) or Alexa Fluor® 647 dye (far red) fluorochromes (Molecular Probes) were used at 1:500 dilutions. After 2h of incubation at 37°C for 1hr followed by washing steps, cells were mounted with Vectashield with DAPI (Vector Lab, Inc) and observed under a Plan Apo VC 60X/1.40 oil or Plan Fluor 10X/0.30 objectives on an inverted Eclipse Ti Nikon microscope equipped with a Nikon Qi1MC or QImaging-Rolera EMC^2^ camera for image capture. Few images were taken also on Zeiss Confocal Imager LSM800.

For live cell analysis 250 ng of GFP-Ago2 or GFP-Dcp1a encoding plasmid was used for transfection of cells grown in a six well plate. For FRAP experiments, cells were photo bleached, and recovery was monitored for a total duration of less than 4 min. Glass bottom petridishes pre-coated with gelatin were used for cell growth for live cell imaging. Cells were observed with a 60X/N.A.1.42 Plan Apo N objective. Images were captured with a IXON3 EMCCD camera in an ANDOR Spinning Disc Confocal Imaging System on an Olympus IX81 inverted microscope. All images were captured on Nikon Eclipse Ti microscope or ANDOR spinning disc microscope was processed with Nikon NIS ELEMENT AR 3.1 software. P-Body tracking and velocity calculations were performed with IMARISx64 software developed by BITPLANE AG Scientific software. All velocity was calibrated against the net velocity of cell boundary. In FRAP experiments intensities of the photobleached regions were recorded. The net intensity value at each time interval was normalized to the intensity of the same region before photobleaching and expressed as a percentage of the initial intensity for plotting. The intensity at the time of photobleaching is set to 0 for comparison of different sets of data.

### RNA isolation and Real time PCR

RNA was extracted with Trizol (Invitrogen) as per manufacturers protocol followed by DNase I treatment (Invitrogen) to remove residual DNA contamination. Real time analyses by two-step RT-PCR was performed for quantification of miRNA on a 7500 REAL TIME PCR SYSTEM (Applied Biosystems) or Bio-Rad CFX96^TM^ real time system using Applied Biosystems Taqman chemistry based miRNA assay system.

miRNA assays were performed using specific primers for human let-7a (assay ID 000377), human miR-122 (assay ID 000445), human miR-21 (assay ID 000397).U6 snRNA (assay ID 001973) was used as an endogenous control. One third of the reverse transcription mix was subjected to PCR amplification with TaqMan® Universal PCR Master Mix No AmpErase (Applied Biosystems) and the respective TaqMan® reagents for target miRNA. Samples were analyzed in triplicates from minimum three biological replicates. The comparative C_t_ method which typically included normalization by the U6 snRNA for each sample was used for all instances. For mRNA quantification, Eurogentec Reverse Transcriptase Core Kit was used to prepare cDNA from RNA sample. Real-time (reverse transcriptase) PCR from cDNA was done with the Mesa Green qPCR Mastermix Plus for SYBR Assay-Low ROX (Eurogentec). 18s rRNA has been used as endogenous control.

### Western Blotting

Western analyses of different miRNP components (Ago2, RCK/p54, and XRN1) were performed as described previously. Detailed list of antibodies used are available in the Information. Imaging of all western blots was performed using an UVP BioImager 600 system equipped with VisionWorks Life Science software (UVP) V6.80.

### Microsome isolation

For microsome isolation, cells were resuspended in 1X hypotonic buffer [10mM HEPES pH 7.8, 1mM EGTA, 25mM KCl] equivalent to three times the packed cell volume (PCV) and incubated for 20min on ice. The cells were spun down and resuspended in 2 volumes of 1X isotonic buffer [10mM HEPES pH 7.8, 1mM EGTA, 25mM KCl, and 250mM Sucrose] and homogenized manually. Post pre-clearing of cell debris and unlysed cells at 1000xg for 10min, mitochondrial fraction was removed at 12,000xg for 15min. The post-mitochondrial supernatant was incubated for 15min with 8mM CaCl2 followed by centrifugation at 8,000xg for 10min to isolate the microsomal pellet. The detailed protocol is described elsewhere [5]. The isolation of polysomes from microsomes by KCl/Puromycin treatment was performed with protocol described elsewhere [4, 5].

### Opti-Prep density grandient based fractionation of cellular organelles

For cell fractionation in iodixanol gradient (OptiPrep® gradient), roughly 2×10^7^ cells were used. The cell pellet incubated in a hypotonic buffer [50mM HEPES pH 7.8, 78mM KCl, 4mM MgCl2, 8.4mM CaCl2, 10mM EGTA, 250mM sucrose, 100μg/ml CHX, 5mM vanadyl ribonucleoside complex, and 1X EDTA-free protease inhibitor cocktail] was homogenized by 30-40 strokes in a glass Dounce homogenizer (Sartorius). The cell homogenate was cleared by centrifugation at 1,000xg twice, and the cleared supernatant was loaded on a 3 to 30% iodixanol gradient (OptiPrep) and ultracentrifuged for 5h at 36,000 rpm in an SW60 rotor (Beckman Coulter). Fractions were collected manually through aspiration and RNA or proteins were isolated for further analysis.

### Sucrose density gradient based fractionation of polysomes

For polysome analysis, approximately 2×10^7^ cells, grown to the desired level of confluency, were lysed in a buffer containing 10mM HEPES pH 8.0, 25mM KCl, 5mM MgCl2, 1mM DTT, 5mM vanadyl ribonucleoside complex, 1% Triton X-100, 1% sodium deoxycholate and 1Χ EDTA-free protease inhibitor cocktail (Roche) supplemented with 100 μg/ml cycloheximide. Polysome profiles were obtained by measuring the absorbance at 254nm using ISCO UA-6 absorbance monitor and fractions were collected on ISCO gradient fractionator. RNA and proteins were isolated from each individual fraction and analyzed.

### Single-step Polysome isolation

For one step polysome isolation cell were lysed in a buffer containing 10mM HEPES, pH 8.0, 25mM KCl, 5mM MgCl2, 1mM DTT, 5mM vanadyl ribonucleoside complex, 1% Triton X-100, 1% sodium deoxycholate, and 1Χ EDTA-free protease inhibitor cocktail (Roche) supplemented with 100 μg/ml cycloheximide. Lysates were cleared by spinning at 3,000xg followed by 20,000xg for 10min each and supernatant were loaded on a 30% sucrose cushion and spun for 1h at 31,200 r.p.m in SW60 rotor (Beckman Coulter). The non-polysomal supernatant was removed and polysomes collected below the sucrose layer was diluted with the hypotonic buffer [10mM HEPES pH 7.8, 25mM KCl, 5mM MgCl2, 1mM DTT] and spun for another 30min at the same speed. The supernatant was removed and polysome pellet was re-suspended in hypotonic buffer mentioned above.

### Flow Cytometry Based Mitochondrial Membrane Potential estimation

JC-1 Dye (Life Technologies), a mitochondrial membrane potential sensitive probe (Life Technologies) was used to label cells according to manufacturer’s protocol. Cells were harvested and incubated with 1mM of JC-1 in the culture medium for 15 mins at 37°C in presence of 5% CO2. Cells were then harvested and washed with 1X PBS and analyzed on a BD FACS CALIBUR instrument using the manufacturer’s protocol.

### Measurement of Mitochondrial Oxygen Consumption

HeLa and MDA-MB-231 cells were plated accordingly as low (50% confluency) and high (100% confluency) density cultures in 24 well cell plates (Seahorse bioscience) in DMEM growth medium containing 10% FBS and placed in a 5% CO2 incubator. On the following day, DMEM base medium (Seahorse Bioscience, North Billerica, MA, USA) was supplemented with 25μM D-glucose and 1μM sodium pyruvate (adjusted to pH 7.4) was added to the cells and incubated for 1h in a non-CO2 incubator at 37°C. The three injections ports (A-C) of the XFe cartridge were loaded with Oligomycin (Oligo, 1μM), Carbonyl cyanide-4-trifluoromethoxyphenylhydrazone (FCCP, 0.5 μM) and Rotenone and antimycin A (Rot + Anti A, 1μM) respectively followed by equilibration and calibration in the instrument for 12 min. Following this, the cell plate was loaded to initiate measurement of oxygen consumption rate (OCR) following the 3min wait, 2min mix and 3min measure cycle over a total period of approx. 1hr. Protein estimation was done by ELISA using Bradford’s reagent to normalize the obtained OCR values OCR rate which was expressed in pmol/min/mg protein.

### *In vitro* RISC cleavage assay

*In vitro* RISC cleavage assay with affinity-purified miRISC-122 was carried out using a 36 nt RNA **5’-AAAUUCAAACACCAUUGUCACACUCCACCAGAUUAA-3’** bearing the sequence complementary to mature miR-122. Reaction was carried out in a total volume of 30 μl with 10fmol of 5’ γ^32^P-labelled RNA in buffer [100mM KCl, 5.75mM MgCl2, 2.5mM ATP, 0.5mM GTP] and protein equivalent amount of RISC at 30°C for 30min. After RNA isolation, products were electrophoresed on a 12% denaturing 8M Urea–PAGE and visualized by autoradiography.

### Exosome Purification and Treatments

The standard exosome purification procedure was based on differential ultracentrifugation. The first steps were designed to eliminate large dead cells and large cell debris by successive centrifugations at increasing speeds. The cleared, conditioned medium was centrifuged for 20 min at 2,000×g, 4°C. Subsequently the supernatant was pipetted off and centrifuged for 30 min at 10,000×g, 4°C. Careful removal of all the supernatant was ensured such that none of the pellet contaminates the supernatant. The residual supernatant is subjected to a single filtration step using a 0.22 μm filter. This eliminated any residual dead cells and large debris while keeping small membranes intact for further purification by ultracentrifugation. The collected supernatant was gently layered over a 2M sucrose cushion and subsequently centrifuged for at least 70 min at 100,000×g, 4°C. Post ultracentrifugation the supernatant over the sucrose cushion level is gently removed by aspiration. The remaining volume containing exosomes is diluted in 1X PBS and centrifuged for at least 30 min at 100,000×g, 4°C. The pellet obtained was resuspended in each tube in 1 ml PBS, using a micropipette. The resuspended pellet in PBS was centrifuged for 1h at 100,000×g, 4°C. To resuspend the final pellet (i.e., exosomes) a small volume of PBS or PLB was added. To avoid contamination from exosomes present in serum used in the culture medium, exosome pre-cleared serum was used to grow the cells for exosome measurement experiments. The protocols were followed as they are described earlier [28, 29, 54].

### Post imaging analysis & others

All western blot images were processed with Adobe Photoshop CS4 for all linear adjustments and cropping. All images captured on Nikon Eclipse Ti microscope or ANDOR spinning disc microscope were processed with Nikon NIS ELEMENT AR 3.1 software. P Body tracking and velocity calculations were performed with IMARISx64 software developed by BITPLANE AG Scientific software. Image cropping was done using Adobe Photoshop CS4. All graphs and statistical analyses were generated in GraphPad Prism 5.00 (GraphPad, San Diego, CA). Two sample Student’s t test was used for analysis. P values < 0.05 were considered to be statistically significant and > 0.05 were not significant (ns). Error bars indicate mean ± SEM.

## Acknowledgement

We acknowledge Witold Filipowicz for different plasmids constructs and also for valuable discussions. SNB is supported by The Swarnajayanti Fellowship from Dept. of Science and Technology, Govt. of India, while YC, SB, SC and SG received their support from CSIR, India. We were supported by funds from High Risk High Reward Project Grant, Dept. of Science and Technology, Govt. of India.

## Author contributions

S.N.B designed research and analyzed data; S.C., Y.C., S.B. and S.G. performed research and analyzed data; S.N.B., S.C. and Y.C. wrote the paper.

The authors declare no competing interest.

## Supplementary Information

### Inventory of Supplementary Information

The Supplemental Information file contains five supplementary figures, one supplementary table and four supplementary videos.

### Supplementary Figure legends

**Figure S1.**
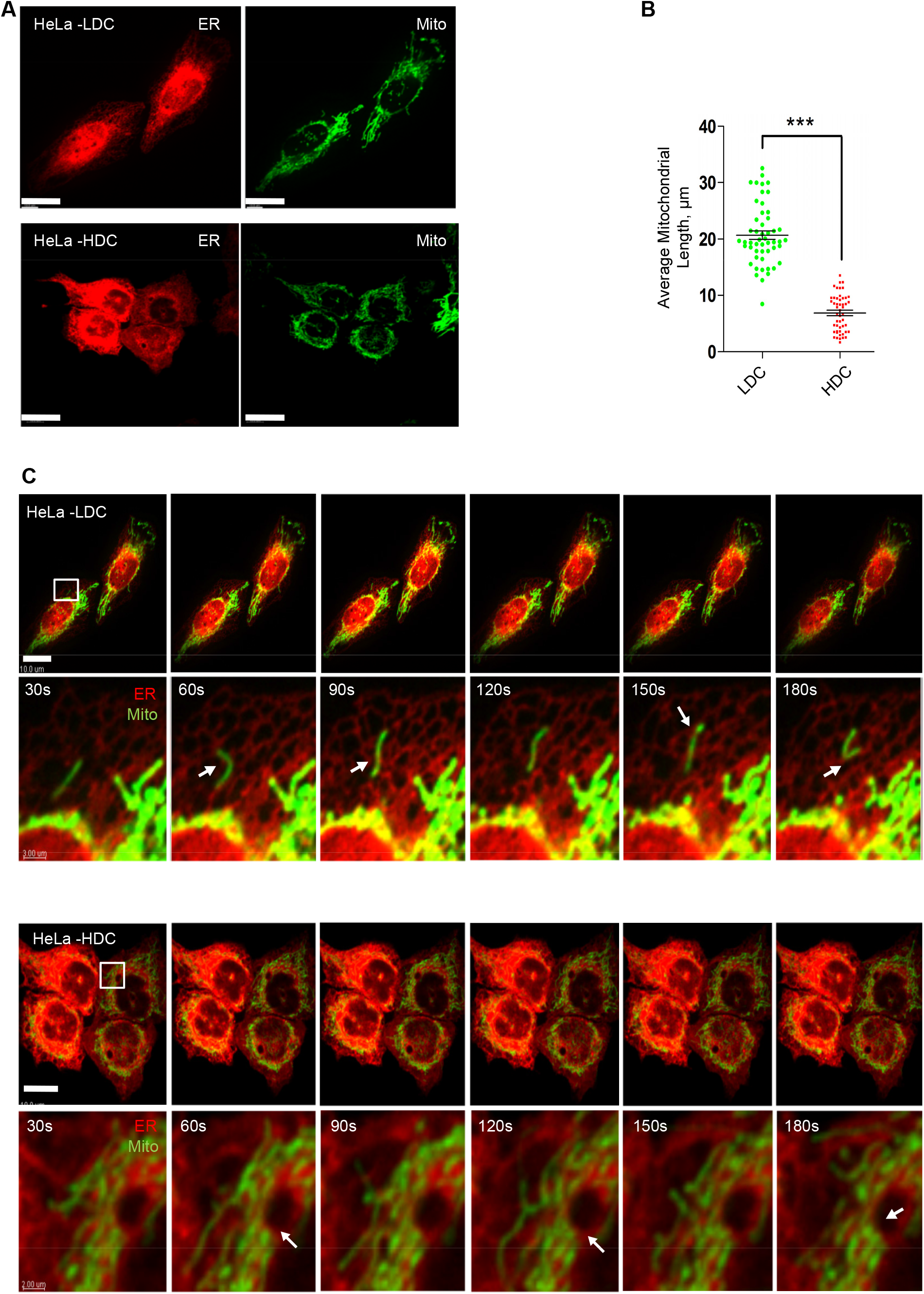
Altered mitochondrial shape and dynamics in HDC state HeLa cells. **(A)** Different mitochondrial shapes of mitochondria in HDC and LDC HeLa cells. A representative picture of cells labeled for ER and mitochondria in HDC or LDC state HeLa cells. Cells expressing a mitochondrial targeting variant of GFP (MT-GFP, green) and an ER targeting variant of DsRed (pDsRed2-ER, red) were used. Bar lengths are of 10μm. (**B)** Mitochondrial size distribution in LDC or HDC state HeLa cells. Quantification based on the images taken for cells in HDC and LDC states. (**C)** Live cell microscopy was done for a total of 3 mins at 1 fps. Representative frames of ER, EE and mitochondria in LDC or HDC HeLa cells for depicted time periods are shown. Cells expressing a mitochondrial targeting variant of GFP (MT-GFP, green) and an ER targeting variant of DsRed (pDsRed2-ER, red) were used. Bars measure 10μm. The ROIs as depicted are 10X zoomed. Also see Supplementary Video 3 and 4. Shown are the mean and s.e.m for n>10. P values are calculated by Student’s t test, and one, two, and three asterisks represent p values less than 0.05, 0.01, and 0.001, respectively.

**Figure S2.**
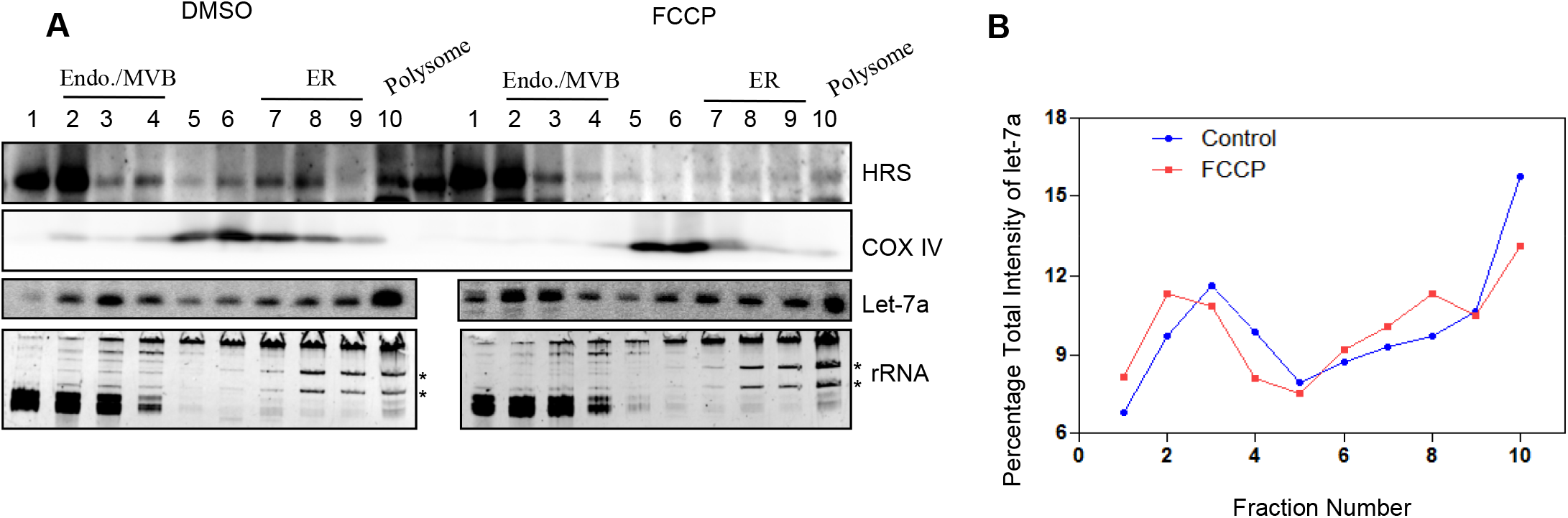
Distribution of miRNP with different subcellular fractions in FCCP treated mammalian cells. **(A)** Representative OptiPrep® gradient (3–30%) analysis of HeLa extracts of indicated treatment by western blotting of specified proteins done with individual fractions. rRNA positions are marked by * on a RNA profile picture done for each fractions. (**B)** Levels of miRNA let-7a (quantified by densitometry) in individual fractions of the Optiprep^®^ from *panel* ***A*** were plotted. Let-7a levels in every individual fraction have been normalized against the total of let-7a band intensity present in all the fractions. The resultant values were fitted on a scale of 100 where “100” depicts the total sum of let-7a density for all the fractions of an individual experiment. p values are calculated by Student’s t test, and one, two, and three asterisks represent p values less than 0.05, 0.01, and 0.001, respectively. Shown are the mean and s.e.m for n>3.

**Figure S3.**
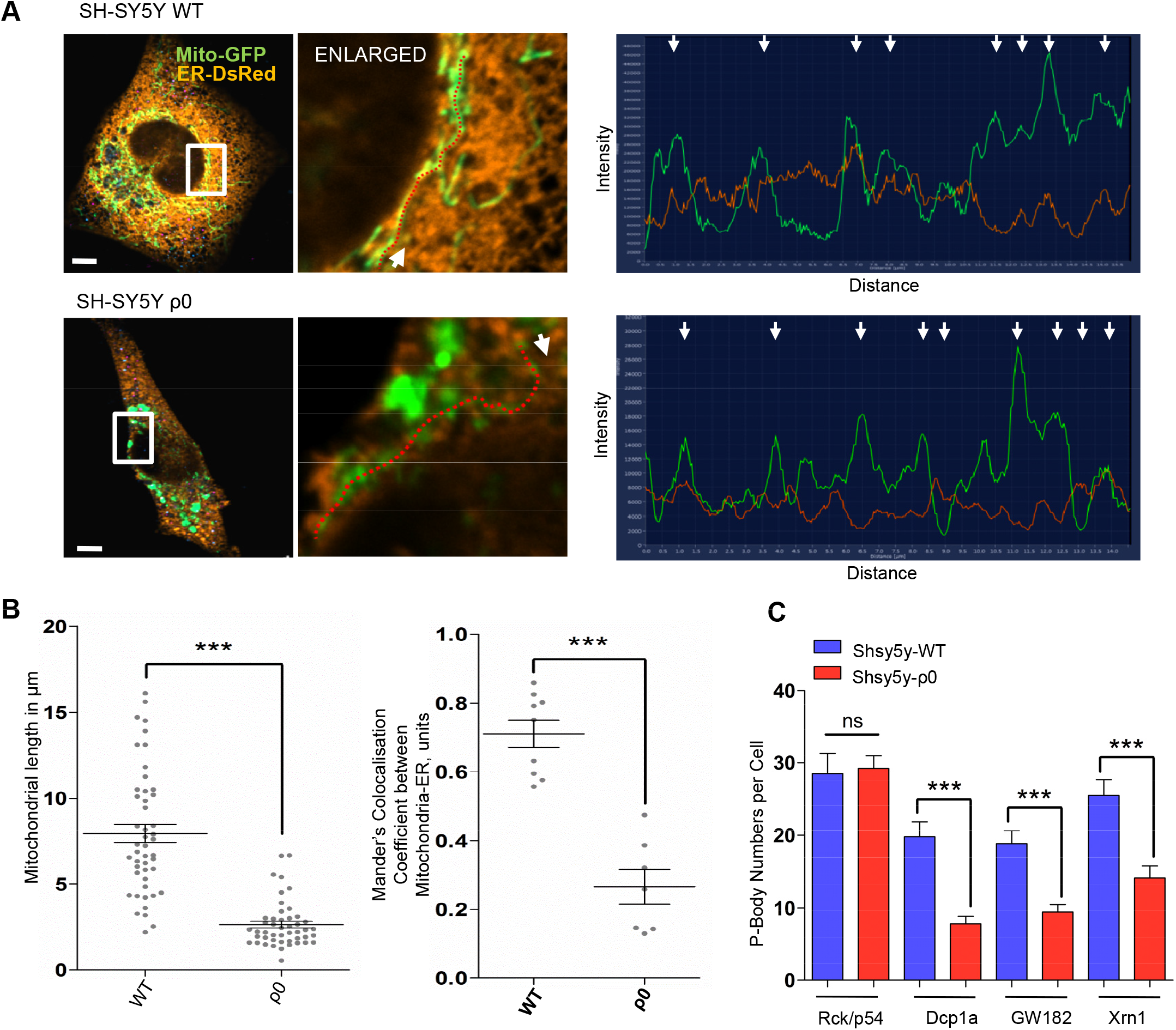
Reduced P-body number, mitochondrial length and mitochondria-ER colocalization in SH-SY5Y ρ0 cells. **(A)** Representative confocal microscopy images (*left panel*) depicting mitochondria (*green*) and ER (*orange*) in WT or ρ0 SH-SY5Y cells were used to determine the line intensity profile (*right panel*) along an arbitrary linear path (*as depicted by dotted red line, left panel*) whose starting direction has been shown by a white arrow within the selected ROI. Arrows in the intensity profile indicate different areas of varied intensity overlap. Cells co-expressing a mitochondrial targeting variant of GFP (MT-GFP, green) and an ER targeting variant of DsRed (ER-DsRed, orange) were used in the merged panels of WT or ρ0 SH-SY5Y cells in *panel **A**.* **(B)** Left panel, mitochondrial length quantification was done for WT or ρ0 SH-SY5Y cells co-expressing a mitochondrial targeting variant of GFP (MT-GFP) using Imaris for n>25 cells. Right panel, mitochondria-ER colocalization was estimated using Mander’s colocalization coefficient based quantification between MT-GFP and ER-DsRed for WT or ρ0 SH-SY5Y cells for n>30 cells. (**C)** Estimation of endogenous Dcp1a, GW182, Rck/p54 and Xrn1 positive body numbers were determined from confocal microscopy frames of WT or ρ0 SH-SY5Y cells labelled for the individual p-body markers by immunofluorescence and normalized against number of cells for n>20 cells. P values are calculated by Student’s t test, and one, two, and three asterisks represent p values less than 0.05, 0.01, and 0.001, respectively whereas ns depicts p values greater than 0.05. Shown are the mean and s.e.m from at least three independent experiments.

**Figure S4.**
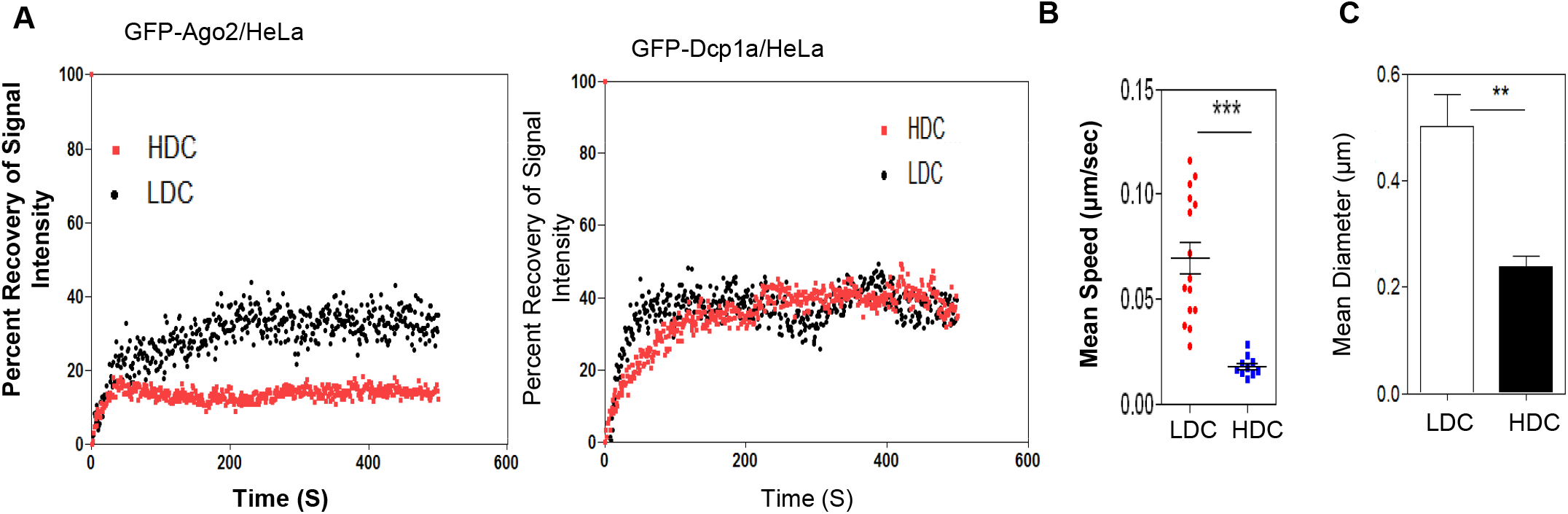
Characteristics of P-Bodies changes with cell confluence. **(A)** FRAP analysis from 100 cells (n>4) transfected with a GFP tagged variant of Ago2 or Dcp1a for indicated experimental sets was done. GFP tagged proteins was photobleached and Images were acquired at 1 frame per second for 500 seconds. The mean percent intensity values of different regions of interest (ROI) thus obtained was plotted. (**B)** Cells transfected with a GFP tagged variant of Ago2 (Ago2-GFP, green) for indicated experimental sets were used. Ago2-GFP live cell images were acquired at 1 frame per second for 500 seconds. Particle Tracking was used to determine the path and speed of the Ago2-GFP bodies. (**C)** Cells transfected with a GFP tagged variant of Ago2 (Ago2-GFP, green) for indicated experimental sets were used. Ago2-GFP fixed cell images were used for calculating the diameter of Ago2-GFP bodies in HeLa HDC or LDC cells.

**Figure S5.**
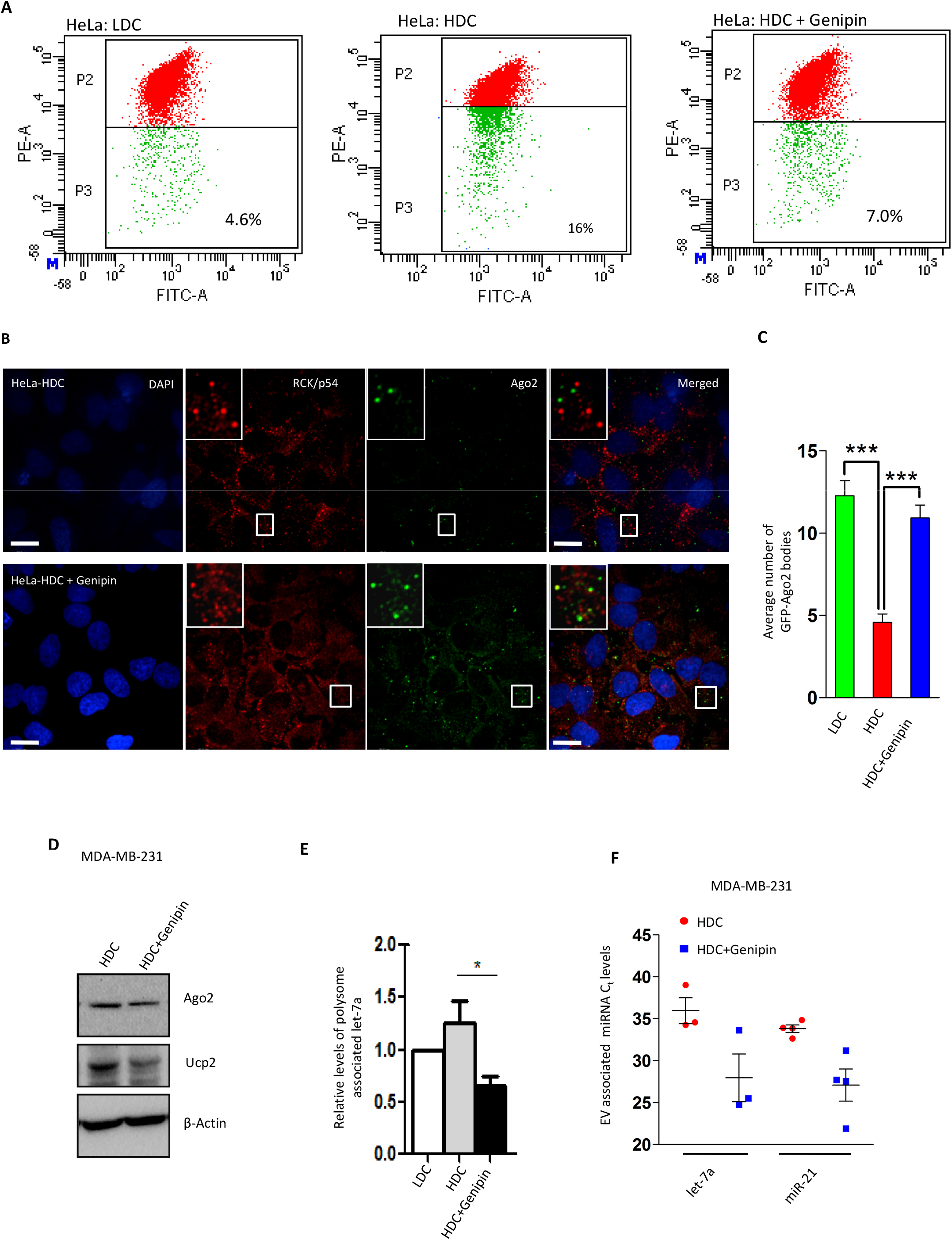
Restoration of mitochondrial membrane potential rescues miRNP trafficking and P-bodies. **(A)** Representative plots using flow cytometry based quantification of JC-1 staining of LDC, HDC and 3 hrs of 100μM genipin treated HDC HeLa cells. Percent of cells in P3 zone (low JC-1 Fluorescence, green) were measured and shown as depicted. (**B)** Representative confocal microscopy images depicting P-bodies in HDC HeLa and HDC HeLa cells treated with Genipin. Indirect immunofluorescence was used to detect endogenous Ago2 (*green*) and RCK/p54 (*red*). Bars measure 10μm. DAPI was used for depicting nuclei. The ROIs as depicted are zoomed 5 times. (**C)** Estimation of numbers of Ago2-positive bodies was done for HDC HeLa or HDC HeLa cells treated with Genipin as shown in the representative images in analysis of indicated proteins. (**D)** The effect of Genipin on Ago2 and Ucp2 protein levels in HDC MDA-MB-231 cells. Representative western blots are shown. (**E)** qRT-PCR based relative quantification of endogenous miR-21 levels in fractions obtained from one-step polysome separation gradients done with HDC or LDC MDA-MB-231 cells of indicated Genipin treatment. Values were normalized against respective miRNA levels in LDC cells and shown in the right panel here. Shown are the mean and s.e.m from at least three independent experiments. (**F)** qRT-PCR based quantification of changes in EV-associated miRNA levels following exposure to Genipin. C_t_ values of endogenous miRNA levels in HDC or LDC MDA-MB-231 cells after Genipin treatment. p values are calculated by Student’s t test, and one, two, and three asterisks represent p values less than 0.05, 0.01, and 0.001, respectively.

**Video 1** Mitochondrial movements in MDA-MB-231 LDC cells.

Time lapse imaging of Mitochondria and ER in MDA-MB-231 LDC cells imaged for 1 frame per second for 115 frames. Cells have been co-transfected with pDsRed2-ER (*red*) and pAcGFP1-Mito (*green*) plasmids. The video plays at 5 frames per second. Scale bars, 10 μm. Time is in hours:minutes:seconds. Also see Figure 1C.

**Video 2** Mitochondrial movements in MDA-MB-231 HDC cells.

Time lapse imaging of Mitochondria and ER in MDA-MB-231 HDC cells imaged for 1 frame per second for 115 frames. Cells have been co-transfected with pDsRed2-ER (*red*) and pAcGFP1-Mito (*green*) plasmids. The video plays at 5 frames per second. Scale bars, 10 μm. Time is in hours:minutes:seconds. Also see Figure 1C.

**Video 3** Mitochondrial movements in HeLa LDC cells.

Time lapse imaging of Mitochondria and ER in HeLa LDC cells imaged for 1 frame per second for 115 frames. Cells have been co-transfected with pDsRed2-ER (*red*) and pAcGFP1-Mito (*green*) plasmids. The video plays at 5 frames per second. Scale bars, 10 μm. Time is in hours:minutes:seconds. Also see Supplementary Figure 1C.

**Video 4** Mitochondrial movements in HeLa HDC cells.

Time lapse imaging of Mitochondria and ER in HeLa HDC cells imaged for 1 frame per second for 115 frames. Cells have been co-transfected with pDsRed2-ER (*red*) and pAcGFP1-Mito (*green*) plasmids. The video plays at 5 frames per second. Scale bars, 10 μm. Time is in hours:minutes:seconds. Also see Supplementary Figure 1C.

**Table S1.** List of Antibodies

| **Name of Antigen** | **Raised in** | **Dilution (WB)** | **Dilution (IF)** | **Source** |
| --- | --- | --- | --- | --- |
| Ago2 (eIF2C2) | Mouse | 1:1000 | 1:100 | Abnova |
| HA | Rat | 1:1000 | 1:100 | Roche |
| HRS | Rabbit | 1:2000 | - | Bethyl |
| Calnexin | Rabbit | 1:10000 | - | Bethyl |
| β-Actin | HRP-conjugated | 1:10000 | - | Sigma Aldrich |
| GW182 | Rabbit | - | 1:400 | Bethyl |
| Alix | Mouse | 1:100 | - | Santa Cruz |
| CD-63 | Mouse | 1:500 | - | BD Biosciences |
| DDX6 /RCK/p54 | Rabbit | 1:10000 | 1:1000 | Bethyl |
| Rab5 | Rabbit | - | 1:100 | Cell Signalling |
| Grp-78/Bip-1 | Rabbit | 1:2000 | - | Millipore |
| Cox-IV | Rabbit | 1:1000 | - | Cell Signalling |
| Ucp2 | Rabbit | 1:4000 | - | Novus |
| XRN1 | Rabbit | 1:10000 | 1:1000 | Bethyl |
| cMyc | Mouse | - | 1:100 | SantaCruz |
| Dcp-1a | Mouse | 1:1000 | 1:100 | Novus |
| VDAC | Rabbit | 1:2000 | - | Sigma Aldrich |
| Ribosomal S3 | Rabbit | 1:1000 | - | Cell Signalling |
| eIF-4E | Rabbit | 1:10000 | 1:1000 | Bethyl |
| P-eIF-4EBP! | Rabbit | 1:500 | - | Cell Signaling |
| P-eIF-2α | Rabbit | 1:500 | - | Cell Signaling |
| eEF2 | Rabbit | 1:10000 | - | Bethyl |
| eIF-2α | Rabbit | 1:500 | - | Cell Signaling |
| mTOR | Rabbit | 1:1000 | - | Cell Signaling |
| P-mTOR | Rabbit | 1:1000 | - | Cell Signaling |
| Ribosomal L7a | Rabbit | 1:1000 |  | Cell Signaling |
| eIF-4E-T | Rabbit | 1:8000 |  | Bethyl |
| Dicer | Rabbit | 1:5000 |  | Bethyl |
| Rictor | Mouse | 1:1000 |  | Sigma |
| β-Tubulin | Mouse | 1:1000 | | SIGMA |

